# Simulation-based assessment of the performance of hierarchical abundance estimators for camera trap surveys

**DOI:** 10.1101/2022.11.23.517687

**Authors:** Bollen Martijn, Casaer Jim, Beenaerts Natalie, Neyens Thomas

**Affiliations:** Centre for Environmental Sciences, UHasselt, Hasselt, Belgium; Research Institute Nature and Forest, Brussels, Belgium; Data Science Institute, UHasselt, Hasselt, Belgium; Leuven Biostatistics and statistical Bioinformatics Centre, KU Leuven, Leuven, Belgium

**Author notes:** **Corresponding author:** Martijn Bollen – Martelarenlaan 42, 3500, Hasselt, Belgium –.

**Keywords:** animal movement, Bayesian inference, count data, Gaussian random walks, population ecology, Stan

## Abstract

The need for knowledge about abundance to guide conservation and management strategies in combination with low detectability of many species has led to a widespread use in ecology and management of a range of hierarchical models (HMs) for abundance. These models also appear like good candidates for inference about local abundance in nature reserves studied by camera traps. However, the best choice among these models is unclear, particularly how they perform in the face of several complicating features of realistic populations that include: (i) movements relative to sites, (ii) multiple detections of unmarked individuals within a single survey, and (iii) low probabilities of detection. We conducted a simulation-based comparison of three HMs (Royle-Nichols, binomial N-mixture and Poisson N-mixture model) in the context of small populations of elusive animals in a single study area, where animals cannot be distinguished individually and hence double counting occurs. We generated count data by simulating camera traps monitoring individuals moving according to a Gaussian random walk. Under the simulated scenarios none of the three HMs yielded accurate abundance estimates. Moreover, the performance of each HM depended on the interpretation of abundance. By pooling abundance estimates for trend estimation, each models’ performance markedly improves. Overall, the Royle-Nichols and Poisson N-mixture models outperform a binomial N-mixture model. This emphasizes the importance of choosing the appropriate HM for the data problem.

## 1 Introduction

Biodiversity monitoring is increasingly important to improve our understanding of factors driving biodiversity changes. To keep up with the need for monitoring ecosystems globally, passive monitoring methods, including camera traps (CTs), have gained popularity (Burton et al., 2015; Delisle et al., 2021). In CT surveys temporally replicated counts are collected at a set of spatial locations, which are informative about animal abundance. Typically, abundance represents a fixed number of individuals at a site for the entire study period (also referred to as season-long abundance; Fogarty and Fleishman, 2021). However, this classical conception of abundance is closely tied to individuals remaining at their site, and abundance models typically do not estimate this quantity well when individuals move across sites. In that case, abundance models actually estimate the frequency of site use, the number of individuals that have used a site at least once (also referred to as superpopulation abundance; Fogarty and Fleishman, 2021). Note that frequency of site use is the equivalence of the ‘proportion of area occupied’ for point abundance surveys in continuous habitats (Efford and Dawson, 2012).

Regardless of its interpretation, true abundances are typically obscured by imperfect detections and possibly also double counting. This requires the use of statistical methods that separate process error, i.e., variability in abundance, from observation error. Over the last decades, a wide variety of hierarchical models (HMs) have been developed for this aim (see Kéry and Royle (2016,2020), for a recent synthesis). In these HMs, the data generating mechanism is represented by a mixture of distributions. The first distribution accounts for variation in the unknown abundance (representing the ecological process). Given the true abundance, the second distribution captures variation in repeated counts obtained from multiple, independent samples of the underlying abundance (representing the detection process; Kéry and Royle (2016)). Two such HMs are the Bernoulli-Poisson mixture model (BernP, also called “Royle-Nichols model”; Royle and Nichols (2003)) and the Binomial-Poisson mixture model (BP, also called “Binomial N-mixture model”; (Royle, 2004)). These HMs are often applied to, respectively, detection/non-detection data and count data when abundance estimates are desired. For parameters to be estimable, these HMs require data to be collected over multiple sites, and repeatedly within a site.

A problem with the use of these models is that they often rely on assumptions that are easily, and commonly, violated. For instance, BP and BernP assume that (i) the number of individuals present at a site does not change over the sampling period, commonly referred to as the closure assumption (note that we will refer to this assumption as “geographical closure” and to populations that satisfy this assumption as being “closed”), (ii) no false-positive detections occur, (iii) detections are independent of each other and (iv) detection probability is equal among all individuals. In practice, these assumptions often do not hold, as animals do not move independently, and they do this throughout a heterogeneous region with boundaries that are often defined based on practical, instead of ecological considerations.

There have been several studies that assess the performance of HMs under the violation of these assumptions (i-iv). While few authors have focused on the BernP so far, the BP model has been evaluated in more depth: Fogarty and Fleishman (2021) show that even small violations of the closure assumption lead to substantial bias in abundance estimated from BP. Moreover, accidental double counting of individuals, which can be viewed as false positive detections, resulted in strong positive biases in this model (Link et al., 2018). To address the occurrence of double counts, Nakashima (2020) fitted a Poisson-Poisson (PP, also called “Poisson N-mixture model”) mixture model, hence accommodating this source of false positives. Martin et al. (2011) induced non-independent detections by simulating from a beta-binomial N-mixture model and found that abundances are overestimated by the BP. We are unaware of a study investigating the effect of unequal detection probabilities among individuals on the estimation quality of the BP model. In addition to assumption violations (i-iv), unmodelled heterogeneity in abundance and detection probability is known to produce biased abundances, particularly when data are sparse (Veech et al., 2016; Duarte et al., 2018; Link et al., 2018). Unmodelled site heterogeneity often arises from erroneously specifying the distribution for the latent state of interest. In many ecological applications, excess zeros may lead to overdispersed counts relative to the Poisson assumption (Martin et al., 2005; Dénes et al., 2015). Modelling excess zeros in these models is possible through negative-binomial or zero-inflated Poisson distributions for the latent abundance states (Wenger and Freeman, 2008). While negative binomial mixtures are often favoured by information criteria, they suffer from what has been called “the good-fit-bad-prediction dilemma”, *i*.*e*., they fit the data well, but lead to ecologically unrealistic parameter estimates. Zero-inflated Poisson models mostly lead to sensible results (Joseph et al., 2009; Kéry and Royle, 2016), making Binomial – zero-inflated Poisson (BZIP) and Poisson – zero-inflated Poisson (PZIP) models suitable candidate HMs for abundance estimation.

The evaluation of HMs remains largely based on data generated by a thinned Poisson point process (but see Neilson et al. (2018); Nakashima (2020) for movement-based data generation). This is an appealing strategy, since the process captured by the model is exactly the same process that generated the data for that model. Hence, consistent failure to recover the true process directly indicates bad model behaviour. In reality, however, detections from CTs emerge by animals moving across a region of interest, and occasionally crossing their detection field. In this study, we simulate detections accordingly by setting up random-walk based movement models, inspired by empirical wild boar movements. We therefore explicitly choose to design a simulation study where observations come from a process that mimics the behaviour of the animals under study, instead of observations that are generated by simulating from a model under investigation. Ultimately, our goal is to compare the estimator quality of BernP, BP, PP, BZIP and PZIP models under various degrees of closure violations as they would emerge in real-life settings from animal behaviour. Specifically, we consider various degrees of (i) closure violations, by changing home range areas relative to a fixed lattice grid; (ii) double count frequency, by changing the population size and movement speed of unmarked individuals; and (iii) detection probability, which is also affected by movement speed. We define estimator quality in terms of bias, root mean square error (RMSE) and 95% confidence interval (CI) coverage in (i) traditional abundance, and (ii) frequency of site use. We anticipate that both (i) and (ii) will be biased, but (i) more strongly than (ii), when closure is violated. Additionally, we expect the PP to outperform the other models overall, given that they accommodate false-positive detections. However, when data are sparse, we hypothesize that models accounting for zero-inflation (BZIP, PZIP) lead to better estimates compared to those that do not account for this. Finally, we also investigate if relative trends in traditional abundances obtained *post-hoc* from HMs are accurate.

## 2 Material and Methods

### 2.1 Simulated space

We simulated a space with a total surface area of 116.64 km^2^ (10.8 km * 10.8 km; **Figure 1**), which roughly equals that of the Hoge Kempen National Park (NPHK) situated in Belgium (Wevers et al., 2020). Next, we simulated a grid layer consisting of 144 grid cells of 0.9 km x 0.9 km, for the placement of camera traps (CTs) (**Figure A.1**). Our grid cells are nine times coarser than the grid cells in Wevers et al. (2020), such that the area (0.81 km^2^) of a single grid is closer to the empirical home range sizes of wild boar (Podgórski et al., 2013; Fattebert et al., 2017). Next, we placed a camera in 25% (36) of the grid cells, according to a randomised regularly spaced sampling design. Within sampled grid cells, a CT “detector” was then simulated by placing a single camera at its centroid with a detection field radius *r* = 15*m* and an angle of view *θ* = 42°, based on camera specifications of the Reconyx Hyperfire HC600 (Reconyx, 2017). The detection fields (“field of views”; FOV) of all cameras were simulated facing North.

**Figure 1:**
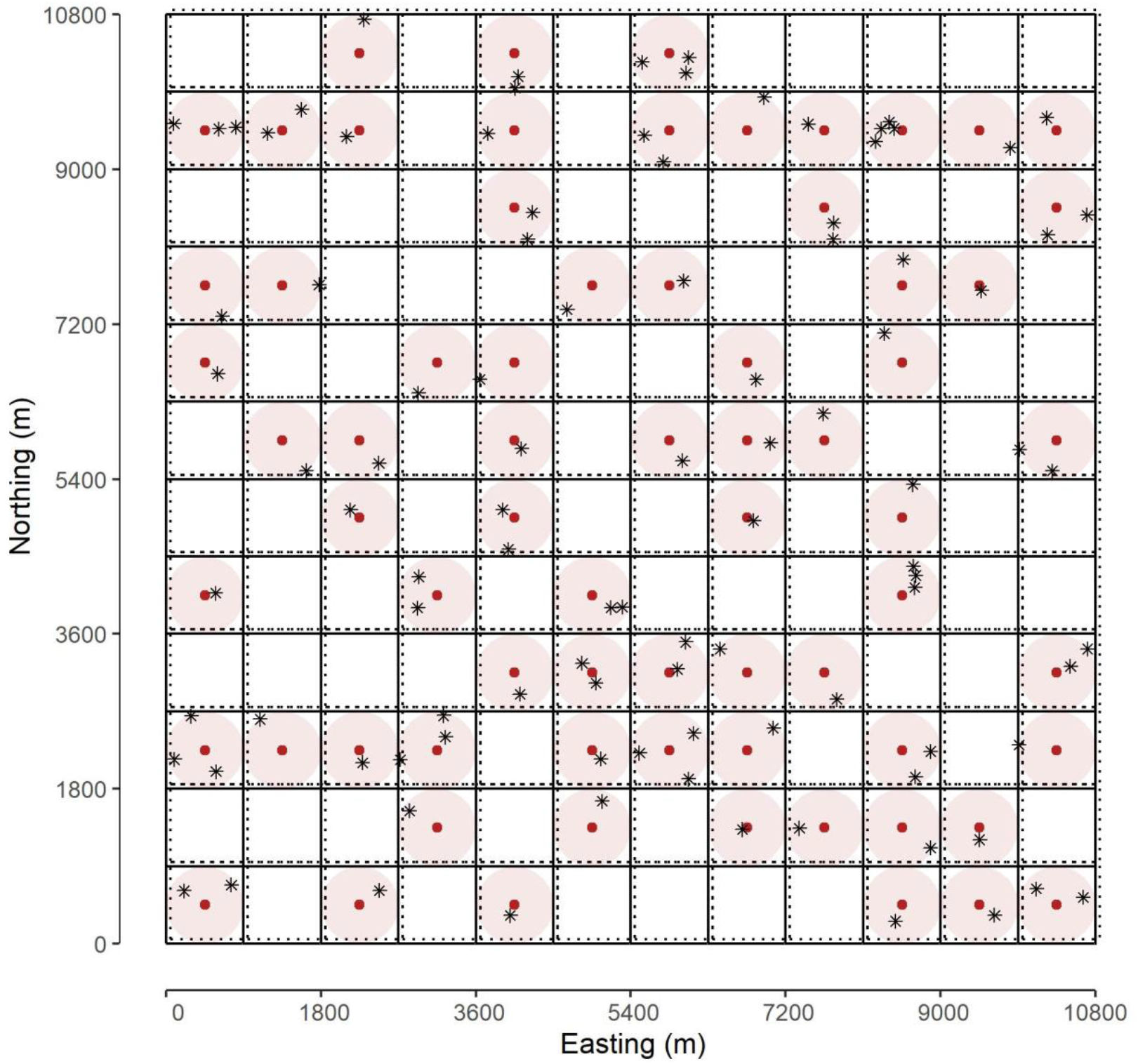
An example of a simulated space. Home range centres (red dots), home range areas (red circles) and the initial positions of individual groups within their home ranges (black asterisks) are indicated. Moreover, sampling cells (solid lines) and bounding boxes for group specific home ranges (dotted lines; shifted slightly for visual clarity) are represented. Note that each home range area may contain more than one group of individuals.

### 2.2 Movement trajectories

Let’s assume that *N* individuals of a species of interest, *e*.*g*., wild boar, live in the region of interest, say *A*. We fix *N a priori* according to **Table 1**, and assume these animals to live in groups. We then simulate forty independent, group-specific and activity-adjusted Gaussian random walks (RWs). To generate realistic animal trajectories for wild boar, but generic enough to apply to a wider range of species, we vary three parameters: the home range area (HRA), the population size (*N*) and the speed of movement, in terms of mean hourly displacement (**Table 1**).

**Table 1:**
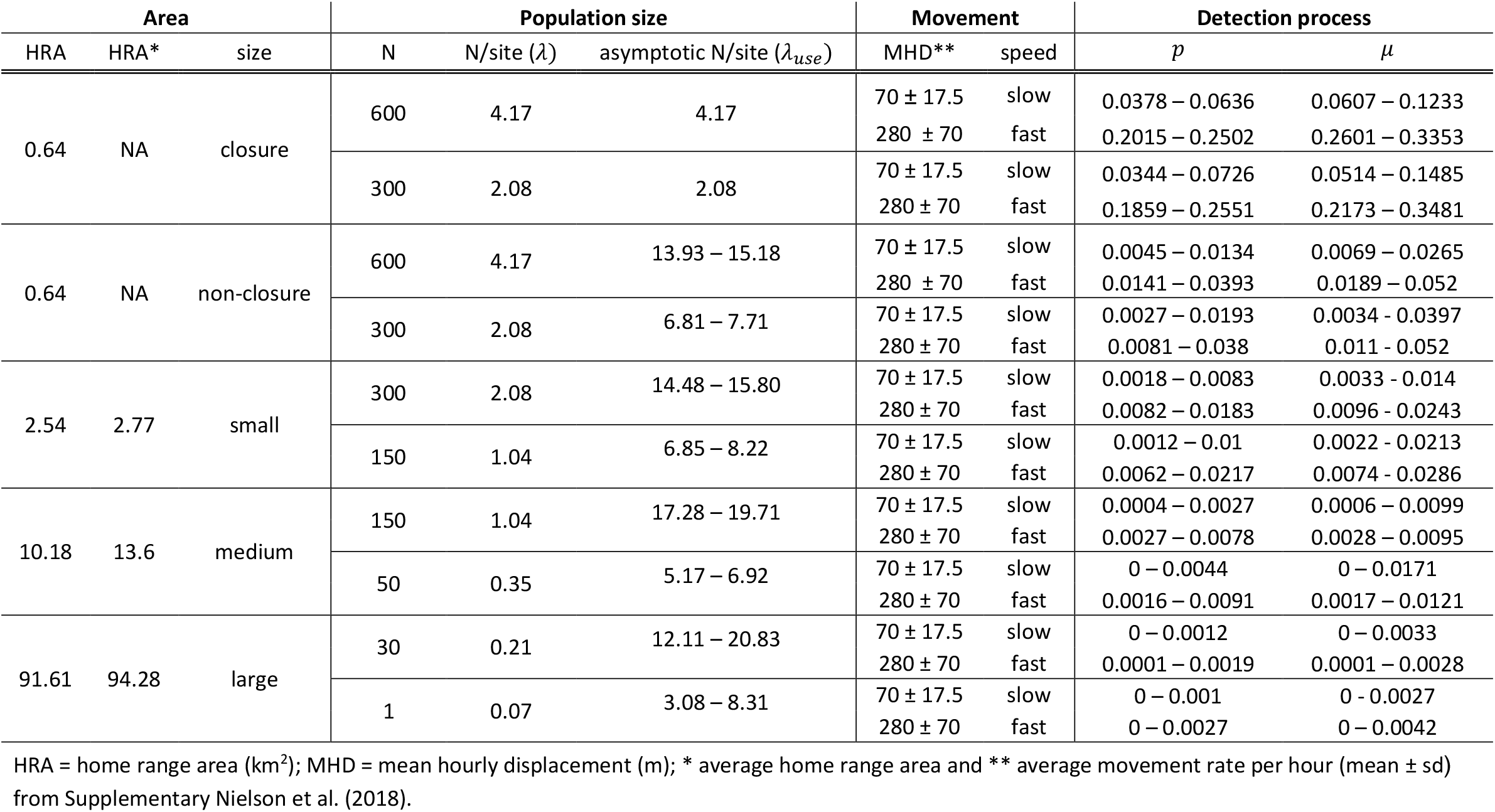
Design of the simulation study, based on three ecological/ behavioural parameters, home range area, population size and mean hourly displacement of (a group of) individual animals. The range of true detection probabilities *p* and rates *μ*, resulting from each scenario, are also indicated.

Let *u* represent a point location, defined by x and y coordinates. For *g* = 1,…, *G* groups of the animal species, *j* = 1, 2,…, 25 days, and *t* = 1,…, *T*, with *T* = 24, hours of the day, the ℝ^2^-valued sequence of vectors ***S*** = ***[X, Y]*** represents steps along x and y vertices. Each *X*_*gjt*_ and *Y*_*gjt*_ is an *i*.*i*.*d*. random variable generated by a univariate normal distribution. However, we discard samples from this distribution, when it results in a position ***u***_*gjt*_ outside of the group-specific HRA, *B*_*g*_. Hence, our RW is constrained to *B*_*g*_, with *B*_*g*_ ⊂ ℝ^2^, and starts at a randomly chosen position ***u***_*g*0_ ∈ *B*_*g*_, such that the vector ***u*** is given by:

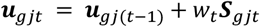

with,

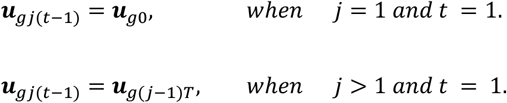

Note that we weigh ***S***_*gjt*_ by an activity level *w*_*t*_. The weighing factor *w*_*t*_ is derived from an activity density over all 24-h periods based on CT data for wild boar obtained from a camera trapping network in the NPHK, using the R-package *activity* (Rowcliffe et al., 2014; Wevers et al., 2020). Moreover, we fix the movement speed by choosing ***S***_*gjt*_ such that the mean hourly displacement and its standard deviation reflect that of **Table 1**. We generated group sizes *N*_*g*_ for each group *g* by sampling empirical group sizes of a wild boar population in the NPHK that satisfy 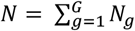. In addition, we simulated 10% and 20% declines in population size for all populations, to explore the ability of the models to estimate relative trends in abundance *post-hoc*. Next, group-specific home ranges *B*_*g*_ were obtained by defining circles with radius *r* and home range centres *c*_*g*_ that are (i) randomly sampled from the entire study space, resulting in closure violations, or (ii) from the fixed set of 0.9 km x 0.9 km grid centroids, resulting in closure. Initial positions of each group ***u***_*g*0_ are randomly positioned within the home range of that group *B*_*g*_, hence ***u***_*g*0_ ∈ *B*_*g*_. Subsequent positions are also bounded by *B*_*g*_, hence ***u***_*gjt*_ ∈ *B*_*g*_, ∀{*g, j, t*}.

When simulating hourly displacements, information on finer scale movements is lost. However, these may be important when they result in the passing or not passing through the FOV (Abolaffio et al., 2019). To ensure that we generate, more realistic, finer scale movements, we simulate Brownian motion between consecutive positions *x* = ***u***_*gj*_(_*t*−1)_ and *y* = ***u***_*gjt*_ using a Brownian Bridge (**Figure A.2**).

### 2.3 Simulating count and detection-non detection data

After obtaining group-specific wild boar trajectories, we generate count data at hourly intervals by assuming that all individuals within one group are detected when they enter the FOV (*i.e., detection probability|trajectory crosses FOV* = 1 for all individuals). Next, we aggregate this information, such that count data are obtained at daily intervals. Since we assumed unmarked individuals, double counts may occur. When only binary inputs are required for particular models, we reduce count data to detection/ non-detection data.

### 2.4 Assessment of criteria for the model inferences

For each simulation replicate, we calculated the true abundance (*λ*), being the mean number of individuals at a grid cell (*i*.*e*., population size *N*/ number of sampled grid cells). This quantity will be used as ground truth, to assess the ability of the HMs to make accurate inferences on *N*. However, the true number of individuals detected, is influenced by both *N* and the HRA of the animal, complicating estimation of *λ*. Hence, we also calculated site use frequency (*λ*_*use*_, *i*.*e*., the mean number of individuals that have used a grid cell at least once during the simulated study period, given that all individuals completely covered their HRA; (Nakashima, 2020; Fogarty and Fleishman, 2021). A graphical summary of sections 2.2 – 2.4 is given in **Figure A.3**.

### 2.5 Statistical models

Within the current study, we assess the performance of HMs, that are commonly used to make inference on abundance. Depending on the data at hand, BernP (for replicated detection/ non-detection data) (Royle and Nichols, 2003) and BP/ PP (for replicated count data) (Royle, 2004; Nakashima, 2020) models are commonly used. For the latter we also tested model parameterisations with a zero-inflated Poisson distribution for the latent abundances (BZIP and PZIP respectively). All these statistical models start from a series of observations *y*_*ij*_ collected at _*i*_ = 1, 2,…, *R* sites, during *j* = 1, 2,…, 25 survey days. Observations *y*_*ij*_ can either consist of binary detection (*y*_*ij*_ = 1)/ non-detection (*y*_*ij*_ = _0_)data or count (*y*_*ij*≥0_)data. The mathematical structure and distributional assumptions for *y*_*ij*_ and *N*_*i*_, the number of individuals present at site _*i*_, are given in **Table 2**. Note that *N*_*i*_ is a latent quantity and its estimation requires marginalisation over *K* support points for the implementation in *Stan*. Here we choose an upper bound *K*, such that *K* = 100 for detection/ non-detection data and *K* =*max*(*y*_*ij*_)+ 100 for count data. Importantly, *p* – the per capita detection probability – represents the probability that an individual is detected (assuming that there are no false-positives), while *μ* – the per capita detection rate – corresponds to the rate with which an individual is detected (when false-positives occur). For notational simplicity, we will refer to these detection parameters jointly as *θ*_*det*_.

**Table 2:**
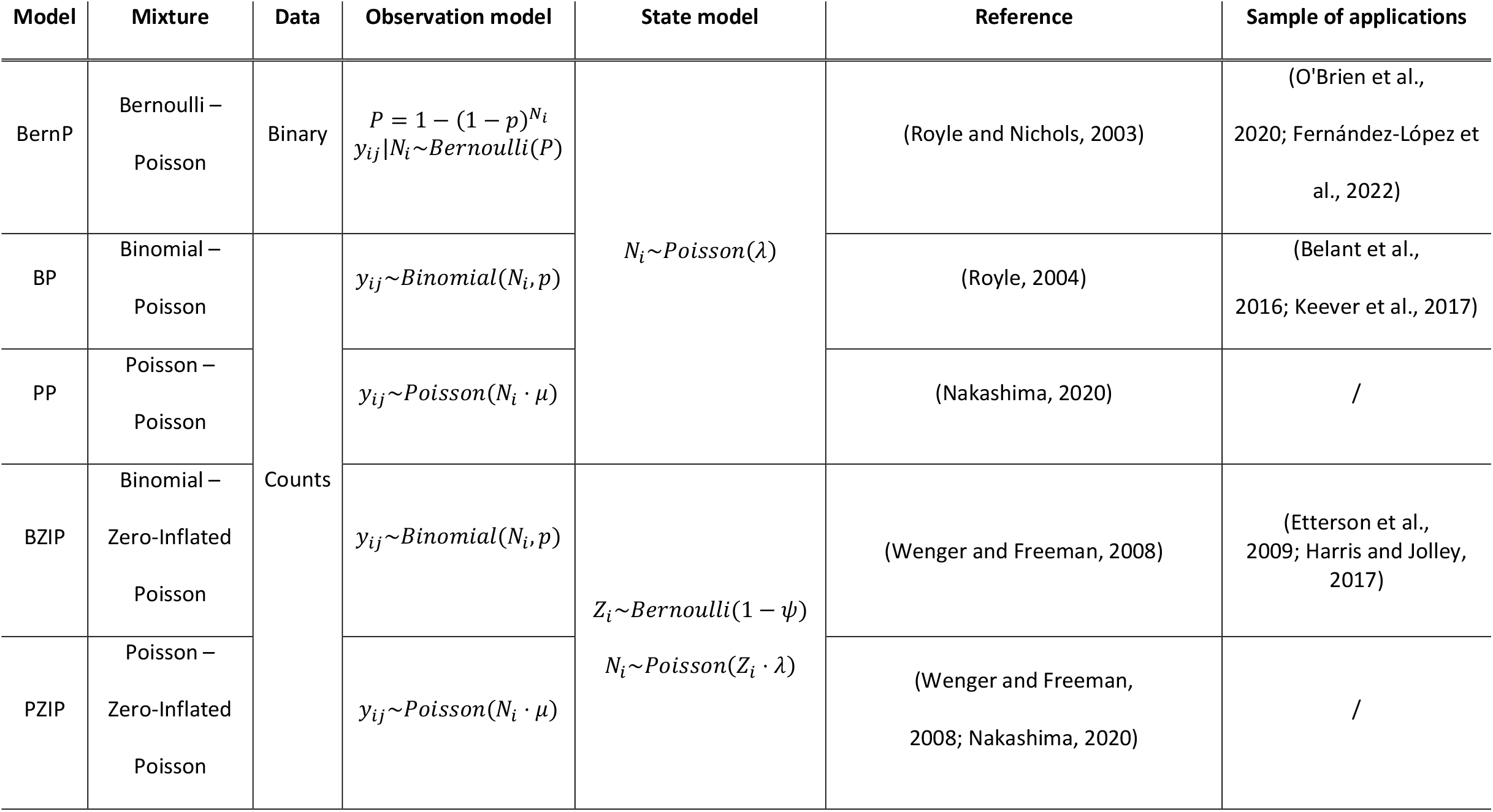
Mathematical structure of the hierarchical models used in this study, their main reference and applications in wildlife research.

Model fitting was performed using *Stan* (via the R package *cmdstanr*), a probabilistic programming language that enables Bayesian estimation through a dynamic Hamiltonian Monte Carlo (HMC) sampler (Carpenter et al., 2017). To assess goodness of fit for competing HMs, we calculate Bayesian *P*-values (Hjort et al., 2006; Link et al., 2018). Additionally, we compute and compare the leave-one-out (LOO) expected log-predictive densities for each model (Vehtari et al., 2021). For more information on the parametrisation, model fitting, goodness-of-fit evaluation and MCMC convergence of HMs in Stan we refer to **Appendix B**. Importantly, we exclude any simulation in which the percentage of divergent chains exceeds 5%, or that results in the potential scale reduction statistic 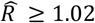 for the parameter of interest from further analysis.

### 2.6 Relative bias, RMSE and Coverage probability

In this study, we will assess estimator performance from the Bayesian HMs of interest, by exploring the bias, relative bias (RB), root mean square error (RMSE) and 95% credible interval (CI) coverage. We calculate the bias as 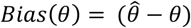; the relative bias as 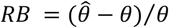; the 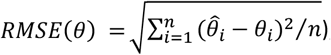and the 95% CI coverage as the proportion of simulations where the true parameter value is enclosed by the 95% CI. We regard 0.95 for this proportion as the threshold value for good uncertainty quantification.

## 3 Results

### 3.1 Goodness-of-fit and predictive performance

The proportions of Bayesian *P*-values *P*_*B*_ and LOO for each candidate model in all simulation scenarios are summarised in **Table B.1**. Overall, the BZIP, PZIP and PP are most frequently identified to have a bad fit, while BernP lead to the best fits according to the *P*_*B*_. Discarding those simulations where *P*_*B*_ < 0.05 did not qualitatively alter our results. On average, the BernP has the highest LOOs, yet for large HRAs, the highest LOOs are observed for BZIP and PZIP.

### 3.2 Accuracy and precision of parameter estimates

We assess estimator quality by inspecting the bias, RMSE and 95% CI coverage for both observation and state parameters for each model under various combinations of study design, population size *N* and HRA. Importantly, we found that increasing the number of grid cells that were sampled beyond 25% had a negligible impact on estimator quality (not communicated). Finally, we omit information about the ZIP models from **Tables 3**-**4**, as they do not result in better estimates for *λ* and *λ*_*use*_ compared to the other models (but see **Appendix C**).

**Table 3:** Summary table for estimator quality of detection parameters *θ*_*det*_, abundances *λ* and site use frequencies *λ*_*use*_ obtained from three Bayesian hierarchical models (BernP, BP, PP). Cells display the proportion of simulation replicates that satisfy |Relative Bias| ≤ 0.5 (note that the highest proportions for each scenario are indicated in bold).

|  |  |  | \|Relative Bias\| ≤ 0.5 |  |  |  |  |  |  |  |  |
| --- | --- | --- | --- | --- | --- | --- | --- | --- | --- | --- | --- |
| | | | Detection parameters $\theta_{det}$ | | | Abundance $\lambda$ | | | Site use frequency $\lambda_{use}$ | | |
| HRA (km²) | N | Speed | BernP | BP | PP | BernP | BP | PP | BernP | BP | PP |
| 0.64<br>(closure) | 600 | slow | **0.73** | 0.10 | 0.35 | **0.15** | 0.00 | 0.13 | **0.15** | 0.00 | 0.13 |
|  |  | fast | **0.98** | 0.08 | 0.05 | **0.35** | 0.00 | 0.05 | **0.35** | 0.00 | 0.05 |
|  | 300 | slow | **0.75** | 0.25 | 0.25 | 0.05 | 0.00 | **0.08** | 0.05 | 0.00 | **0.08** |
|  |  | fast | **1.00** | 0.15 | 0.08 | **0.23** | 0.03 | 0.08 | **0.23** | 0.03 | 0.08 |
| 0.64 | 600 | slow | 0.00 | **0.25** | 0.05 | 0.00 | 0.05 | **0.65** | 0.00 | **0.63** | 0.15 |
|  |  | fast | 0.00 | **0.05** | 0.00 | 0.00 | 0.23 | **0.90** | 0.00 | **0.53** | 0.00 |
|  | 300 | slow | 0.00 | **0.25** | 0.20 | 0.00 | 0.15 | **0.55** | 0.00 | **0.60** | 0.30 |
|  |  | fast | 0.00 | **0.08** | 0.00 | 0.00 | 0.50 | **0.70** | 0.00 | **0.43** | 0.03 |
| 2.54 | 300 | slow | 0.00 | **0.28** | 0.03 | 0.05 | 0.05 | **0.28** | 0.00 | **0.38** | 0.03 |
|  |  | fast | 0.00 | **0.48** | 0.00 | **0.93** | 0.00 | 0.00 | 0.00 | **0.68** | 0.13 |
|  | 150 | slow | 0.00 | **0.30** | 0.13 | 0.05 | 0.00 | **0.10** | 0.00 | **0.40** | 0.13 |
|  |  | fast | 0.00 | **0.63** | 0.13 | **0.88** | 0.00 | 0.03 | 0.00 | **0.45** | 0.30 |
| 10.18 | 150 | slow | 0.00 | **0.05** | 0.03 | 0.00 | 0.05 | **0.18** | 0.00 | **0.45** | 0.03 |
|  |  | fast | 0.00 | **0.40** | 0.00 | **0.88** | 0.00 | 0.00 | 0.00 | **0.58** | 0.00 |
|  | 50 | slow | 0.00 | 0.00 | **0.03** | 0.10 | 0.15 | **0.18** | 0.00 | **0.53** | 0.08 |
|  |  | fast | 0.00 | **0.48** | 0.05 | **0.95** | 0.00 | 0.00 | 0.00 | **0.43** | 0.15 |
| 91.61 | 30 | slow | 0.00 | 0.00 | 0.00 | **0.20** | 0.00 | 0.00 | 0.00 | **0.05** | 0.00 |
|  |  | fast | 0.00 | 0.00 | 0.00 | **0.83** | 0.00 | 0.00 | 0.00 | **0.23** | 0.00 |
|  | 10 | slow | 0.00 | 0.00 | 0.00 | **0.38** | 0.00 | 0.00 | 0.00 | **0.25** | 0.08 |
|  |  | fast | 0.00 | 0.00 | 0.00 | **0.58** | 0.00 | 0.00 | 0.00 | **0.40** | 0.13 |
| Median |  |  | 0.00 | **0.13** | 0.03 | **0.18** | 0.00 | 0.08 | 0.00 | **0.41** | 0.08 |

#### 3.2.1 Detection parameters

For closed populations with HRAs of 0.64 km^2^, only the BernP yields accurate estimates of *θ*_*det*_ across all densities and movement speeds. All other HMs consistently underestimated *θ*_*det*_, particularly for slow-moving populations (**Figure 2a; Table 3**). Additionally, the BernP is the only model that is close to the 95% CI coverage threshold in this situation. When the closure assumption is violated, none of the HMs reaches the CI coverage threshold, despite low RMSEs in the BP and PP-case (**Table C.1**). Moreover, the BernP displays the largest (positive) bias in these scenarios (**Figure 2a**). Overall, 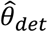 obtained through the BZIP are the most accurate according to the median proportion of simulations with |Relative Bias| ≤ 0.5, followed by the BP and the PP (**Table 3**; **Table C.4**). The BernP displays the strongest absolute bias in *θ*_*det*_ on average.

**Figure 2:**
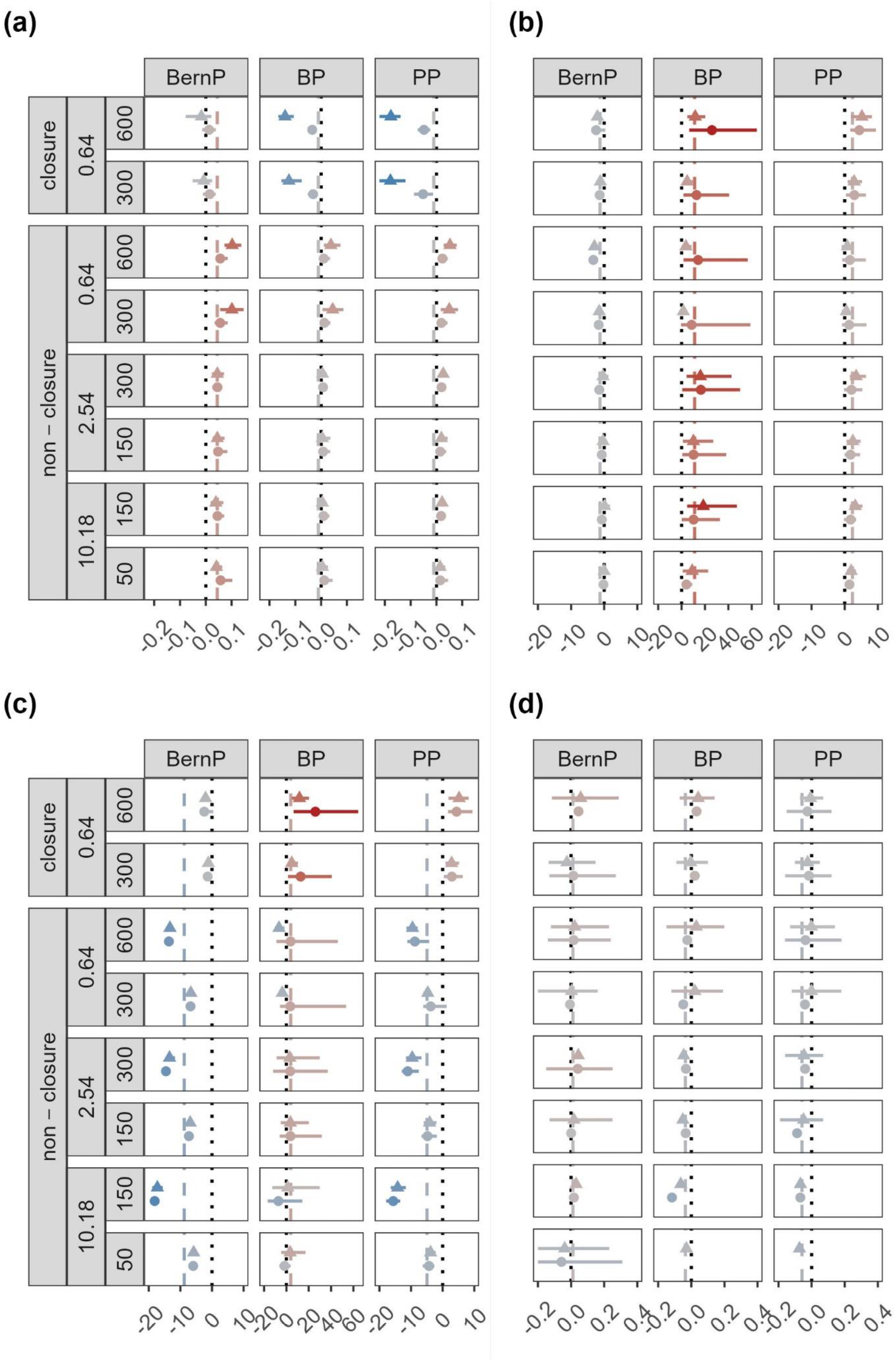
Mean bias (dots/ triangles), together with 2.5% and 97.5% quantiles (solid lines) in the estimated **(a)** detection parameters *θ*_*det*_, **(b)** abundances *λ*, **(c)** site use frequencies *λ*_*use*_ and **(d)** trends in *λ*. Results are displayed for all combinations of population size (N = 10, 30, 50, 150, 300, 600), speed of movement (slow: triangles, fast: dots), home range area (HRA) in km^2^ (0.64, 2.54, 10.18 and 91.81), geographical closure (closure, non-closure) and hierarchical model fitted (BernP, BP, PP, BZIP and PZIP). Line of equality (dotted line). Average bias for each HM (dashed line). For visual clarity, x-scales are different for the subpanels in (b) and (c)

#### 3.2.2 Abundance

Regardless of the simulation scenario, all HMs for count data (*i*.*e*., BP, PP, PZIP and BZIP) consistently overestimate abundance *λ* (**Figure 2b**). Overall, the bias in *λ* is nearly unaffected by the HRA. However, as population sizes *N* are smaller for larger HRA, the relative bias in *λ* strongly increases with HRA. Interestingly, this trend towards inflated estimates of *λ* was much stronger when HMs with a zero-inflated Poisson distribution were fitted for the latent number of individuals (*i*.*e*. BZIP and PZIP). However, for smaller HRAs the PZIP leads to *λ* similar to estimates obtained from PP. Unlike HMs for count data, the BernP only slightly underestimates *λ* in most scenarios (**Figure 2b**). Despite fairly low RMSEs, the BernP was never able to reach the 95% CI coverage threshold (**Table C.2**). According to the median, BernP most accurately estimates *λ*, followed by the PP and BP, with respectively 18%, 8% and 0% of simulations resulting in |Relative Bias| ≤ 0.5 (**Table 3**). Both the PZIP and BZIP perform poorly, due to their tendency to produce inflated estimates of *λ* for large HRAs (**Table C.5**).

#### 3.2.3 Site use frequency

BZIP and PZIP revealed very similar patterns in bias for site use frequency *λ*_*use*_ compared to abundance *λ*. Nonetheless, bias in *λ*_*use*_ for the BernP, PP and BP differs considerably with that observed for *λ* (**Figure 2b-c)**. For small HRAs and closed populations, the PP and BP overestimate *λ*_*use*_, while the BernP very slightly underestimates it. In all other situations, negative bias in *λ*_*use*_ is observed for BernP and PP. Accurate *λ*_*use*_ estimates are obtained from BP for medium HRAs (**Figure 2c**). According to the median, BP most accurately estimates *λ*_*use*_, followed by the PP and BernP, with respectively 41%, 8% and 0% of simulations resulting in |Relative Bias| ≤ 0.5 (**Table 3**). Again, none of the HMs was able to reach the 95% CI coverage threshold (**Table C.3**).

#### 3.2.4 Trend in abundance

When trends in *λ* are of prime interest, PP outperforms both BernP and BP in nearly all scenarios (**Table 4**). Despite some variability in accuracy, **Figure 2d** reveals that the mean estimated trends in *λ* are close to the true trends for all models except the ZIP models, and for all HRAs except 91.61 km^2^. Trends estimated from a BP display higher variability compared to PP and BernP, while BZIP and PZIP consistently underestimate the true trends in *λ*. According to the median, HMs yield respectively 49% and 70% (BernP), 37% and 48% (BP), and 67% and 75% (PP) of trend estimates (10% and 20% decline in population size *N*) with a |Relative Bias| ≤ 0.5 (**Table 4**).

**Table 4:** Summary table for estimator quality of relative abundance (*i*.*e*., trends in abundance estimated by stacking abundance estimates *λ*) obtained from three Bayesian hierarchical models (BernP, BP, PP). Cells display the proportion of simulation replicates that satisfy |Relative Bias| ≤ 0.5 (note that the highest proportions for each scenario are indicated in bold).

|  |  |  | \|Relative Bias\| ≤ 0.5 |  |  |  |  |  |
| --- | --- | --- | --- | --- | --- | --- | --- | --- |
|  |  |  | 10% trend |  |  | 20% trend |  |  |
| HRA (km²) | N | Speed | BernP | BP | PP | BernP | BP | PP |
| 0.64<br>(closure) | 600 | slow | 0.44 | 0.44 | **0.75** | 0.71 | 0.57 | **0.84** |
|  |  | fast | 0.54 | 0.73 | **0.95** | 0.71 | 0.94 | **0.98** |
|  | 300 | slow | 0.66 | 0.39 | **0.69** | 0.74 | 0.50 | **0.77** |
|  |  | fast | 0.71 | 0.86 | **0.93** | 0.89 | 0.96 | **0.98** |
| 0.64 | 600 | slow | 0.63 | 0.48 | **0.75** | 0.83 | 0.53 | **0.80** |
|  |  | fast | 0.72 | 0.77 | **0.87** | 0.86 | 0.88 | **0.93** |
|  | 300 | slow | 0.60 | 0.48 | **0.68** | 0.72 | 0.57 | **0.74** |
|  |  | fast | 0.63 | 0.78 | **0.84** | 0.83 | 0.88 | **0.93** |
| 2.54 | 300 | slow | 0.60 | 0.29 | **0.63** | 0.71 | 0.42 | **0.79** |
|  |  | fast | 0.45 | 0.44 | **0.76** | 0.56 | 0.58 | **0.88** |
|  | 150 | slow | 0.40 | 0.31 | **0.48** | 0.64 | 0.47 | **0.62** |
|  |  | fast | 0.53 | 0.44 | **0.74** | 0.70 | 0.59 | **0.83** |
| 10.18 | 150 | slow | 0.38 | 0.32 | **0.43** | 0.58 | 0.45 | **0.61** |
|  |  | fast | 0.54 | 0.34 | **0.66** | 0.73 | 0.46 | **0.74** |
|  | 50 | slow | 0.17 | **0.27** | **0.27** | 0.34 | **0.35** | 0.23 |
|  |  | fast | 0.41 | 0.32 | **0.50** | **0.62** | 0.42 | 0.58 |
| 91.61 | 30 | slow | 0.09 | 0.14 | **0.18** | **0.30** | 0.26 | 0.17 |
|  |  | fast | 0.25 | 0.27 | **0.38** | 0.46 | **0.47** | 0.41 |
|  | 10 | slow | 0.00 | **0.04** | 0.02 | 0.07 | **0.13** | 0.07 |
|  |  | fast | 0.03 | 0.18 | **0.19** | 0.18 | **0.33** | 0.19 |
| Median |  |  | 0.49 | 0.37 | **0.67** | 0.70 | 0.48 | **0.75** |

## 4 Discussion

Increasing trends in per-capita detectability and detection rate with population size *N*, as indicated in **Table 1**, are within the line of expectations, since denser populations are expected to result in more frequent detection events (*i*.*e*., an individual crossing the FOV). We follow Neilson et al. (2018), in assuming that populations of individuals occupying large HRAs, such as apex predators, are generally less dense (**Table 1**). This assumed confounding of HRA and population size, explains the decreasing trend of *θ*_*det*_ with HRA. Evidently, the per-capita detection probability is consistently lower than the per-capita detection rate, as some individuals may be detected more than once on a survey day. Regardless of geographical closure and the HRA, the abundance *λ* equals *N*/*N*_*site*_, resulting in the densities indicated in **Table 1** (“N/site”). Site use frequency, *λ*_*use*_, is a product of both population size *N* and HRA. In theory, *λ*_*use*_ increases with both increasing *N* and HRA. However, in our simulations these parameters are confounded to simulate realistic animal populations (Neilson et al., 2018). Hence, *λ*_*use*_ is seemingly unaffected by *N* and HRA, as their effects cancel out. Only, the effect of closure violations on *λ*_*use*_ is clearly visible (**Table 1**; “asymptotic N/site”).

In our simulations, except for closed populations occupying HRAs of 0.64 km^2^, true *θ*_*det*_ are extremely small (*i*.*e*., always *θ*_*det*_ < 0.05). Hence, it is perhaps not surprising that in these scenarios positive bias in *θ*_*det*_ persisted across most of the HMs considered. It is notoriously difficult for these HMs to accurately estimate parameters near the boundary of their parameter space (Welsh et al., 2013; Dennis et al., 2015). Thus, we would like to warn practitioners that very small detection probabilities/ detection rates, which are frequent in camera trapping studies, often do not contain sufficient information to confidently analyse within complex modelling frameworks. However, modelling detections of multiple species jointly has the potential to overcome issues related to low detectability (Gomez et al., 2018).

Furthermore, our results reveal that none of the HMs considered in this study were able to estimate *λ* accurately under all scenarios. The BernP model slightly underestimates *λ* in most scenarios as opposed to simulation results of Royle and Nichols (2003). Nonetheless, the BernP model most accurately estimates *λ* on average, despite only using binary input data. A possible explanation for the binary input model to outperform count models lies in the occurrence of double-counting of individuals. Indeed, we assumed that animals were unmarked, and hence could not be individually identified by CTs, which led to badly inflated 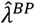, a well-known issue for BP (Duarte et al., 2018; Link et al., 2018). By allowing for false positives, the PP model alleviates a large part of the positive bias in *λ* observed under BP. Nonetheless, 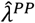 are still positively biased. Specifying a distribution for the latent N_*i*_’s that allows for overdispersion and/ or underdispersion can be a solution to improve inference when extra-variation in counts is present among sites. Here, we used zero-inflated Poisson distribution for N_*i*_’s (Wenger and Freeman, 2008), which did not result in more accurate *λ*’s. On the contrary, both 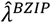and 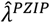 display some of the strongest biases, observed in our simulation study. This is likely a consequence of these models being very complex, *i*.*e*., they require the estimation of two latent parameters in addition to a detection probability/ detection rate, for the sparse data simulated in our study. Despite their inaccurate estimates BZIP and PZIP models were preferred by LOOs, which is possibly because they provide a good description of the many zero-observations present in the data. Negative binomial models have also been suggested to deal with extra-variation in counts, but suffer from a similar “good-fit-bad-prediction dilemma” (Joseph et al., 2009; Kéry and Royle, 2016). Hence, there seems no reason to assume that HMs with negative binomial state models perform any better than the zero-inflated Poisson HMs evaluated here. Another option that we did not explore in this study, but which could improve inference, is the use of a beta-binomial distribution for *p* in the BP case, which can alleviate bias resulting from correlated detections within samples (Martin et al., 2011).

Given the presence of bias in *λ*, the use of HMs for inferring the site use frequency *λ*_*use*_ could be an alternative that provides reliable answers to many wildlife related questions. Site use frequency is effectively a product of population size and HRA, hence reflects both the true number of animals present *N*, as well as their movement pattern (Chandler et al., 2011; Nakashima, 2020; Fogarty and Fleishman, 2021). Our results suggest that HMs do not necessarily estimate *λ*_*use*_ more accurately than *λ*. One important difference, however, is that count models in our study are more suited for inference on *λ*_*use*_ compared to a binary outcome model (*i*.*e*. BernP). This highlights that model choice may depend on the goal of your inference, and specifically the interpretation that is given to state parameters.

Regardless of the capability of HMs to yield unbiased estimates of *λ* (or *λ*_*use*_), it is often more interesting to explore temporal trends in *λ*. Here, we explored trend estimates obtained through comparing abundances *λ* retrieved from two static models *post-hoc* (one fitted to the data before an induced population decline, and the other after the decline). Overall, simulated declines in population size *N* of 10% and 20%, are accurately estimated more frequently than standalone *λ*. However, when a relative bias up to 50% is tolerated, 51%, 63%, 33% and 30%, 52%, 25% of the trends will be misleading for respectively 10% and 20% declines estimated by BernP, BP and PP. This can have profound consequences when decision-making is based on such trend measures.

## 5 Conclusions

We have shown that under realistic movement trajectories, estimating detectability in camera trapping studies will typically be extremely low. In addition, when individuals cannot be individually identified, and thus double counting cannot be excluded, we find strong biases in estimates of abundances *λ* for BP, BZIP and PZIP models. Through accommodating false-positives, the PP model was able to estimate *λ* more accurately. Interestingly, we find the estimator quality of *λ* to be superior for the BernP, while *λ*_*use*_ is more accurately estimated by PP. Finally, we report that focussing on trend estimates, obtained through comparing *λ*’s retrieved from model fits before and after a population decline, yield accurate trend estimates more consistently than standalone *λ*, especially for PP. In conclusion, practitioners should avoid using abundance estimates from single-season HMs and turn to relative trend estimates instead. Depending on the context, model-based approaches, taking into account temporal trends, spatial trends or a combination thereof, might further improve the accuracy of inference from HMs. From this perspective, it would be valuable to assess the estimator quality of temporally and/ or spatially explicit HMs in a simulation study similar to ours. Additionally, it would be interesting to further fine-tune our movement trajectories such that they more closely represent movements of a focal species of choice. Moreover, this would allow researchers to also experiment with design choices, such as camera trap placement, targeted at optimising model performance for the species at hand. Given the multispecies monitoring potential of CTs, it would be interesting to determine to what extent an optimal design, in terms of sampling locations, for one species can be used to make meaningful inference about non-focal species as well.

## Authors’ contributions

MB: methodology, formal analysis, visualisation and writing - original draft preparation.

TN: methodology, validation, and supervision. JC and NB: supervision.

All authors: writing - review & editing, conceptualisation.

All authors contributed critically to the drafts and gave final approval for publication.

## Acknowledgements

MB is a PhD fellow funded by a BOF-mandate at Hasselt University. Services used in this work were provided by the VSC (Flemish Supercomputer Center), funded by the Research Foundation – Flanders (FWO) and the Flemish Government. TN gratefully acknowledges funding by the FWO (grant number G0A4121N) and by the Internal Funds KU Leuven (project number 3M190682). Finally, we would like to express our gratitude towards Marc Kéry for his feedback, which has helped to improve the clarity of this manuscript.

## Conflict of interest

The authors declare no conflict of interest.

## Data accessibility

The datasets generated during and/or analysed during the current study are available on a GitHub repository, https://github.com/MartijnUH/RWsim_abundance_models

## Notes

https://github.com/MartijnUH/RWsim_abundance_models

### Competing Interest Statement

The authors have declared no competing interest.

