## Supplementary material for "Simulation-based assessment of the performance of hierarchical abundance estimators for camera trap surveys": Appendix A_study_design.docx

Appendix A. Visual representation of random walks and study design

The study design described in Section 2 (Materials & Methods) of the main paper, is further visualised in this appendix. For an example of a randomized regularly spaced sampling grid, see **Figure A.1 (a)**. Home range areas of different sizes are visualized in **Figure A.1 (b-f).** Note that the home range areas in subpanels **(b)** and **(c)** are identical. However, in subpanel **(b)** home range centers are sampled from grid centroids, resulting in population closure. In subpanel **(c)** home range centers are sampled randomly from the entire state space.

The movement model explained in Section 2.2 of the main paper, simulates group-specific movement at hourly intervals. However, to make the animal trajectories generated from our model more realistic, we add Brownian motion to each of the line segments representing hourly steps. For example the trajectories displayed in **Figure A.2** **(c-d)** are obtained from **Figure A.2** **(a-b),** after a Brownian bridge, using 6 endpoints (*i.e.*, 10 min. intervals) was added to each set of consecutive point locations $\boldsymbol{u}$. Let $\tau_{t0}=t-1$ and $\tau_{t6}=t$. Then a Brownian bridge starting at $x_{t}$ at time $\tau_{t0}$ and passing through point $y_{t}$ at time $\tau_{t6}$, $\tau_{t6} > \tau_{t0}$, is given by

$$W_{\tau_{t0},x_{t}}^{\tau_{t6}, y_{t}}\left( \tau_{t} \right)=\boldsymbol{x}_{t}+W\left( \tau_{t}-\tau_{t0} \right)-\frac{\tau_{t}-\tau_{t0}}{\tau_{t6}-\tau_{t0}}\cdot\left( W\left( \tau_{t6}-\tau_{t0} \right)-\boldsymbol{y}_{t}+\boldsymbol{x}_{t} \right)$$

And hence the resulting Wiener process at hour $t$ can be written as the vector

$$W_{\tau_{t0}, x_{t}}^{\tau_{t6}, y_{t}}\left( \tau_{t} \right)= [\boldsymbol{u}_{gj\tau_{t0}},\boldsymbol{u}_{gj\tau_{t1}},..., \boldsymbol{u}_{gj\tau_{t6}}]$$

Finally, **Figure A.3** summarizes the entire data generating mechanism (i.e., the realization of the true processes).

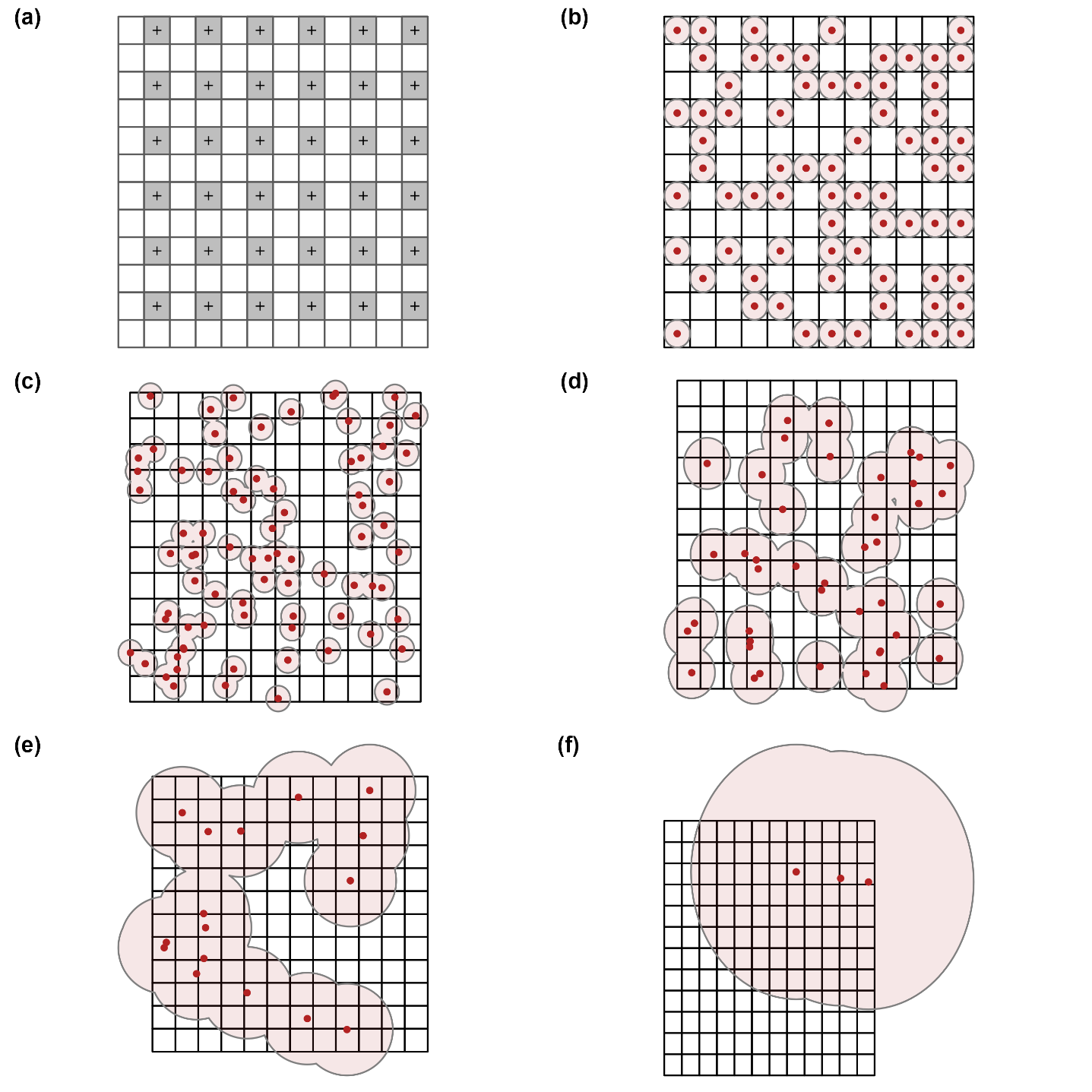

**Figure A.1:** Overview of the sampling design and simulated home range areas. An example of a randomised regular sampling grid for camera placement with a spacing of 0.9 km (a). Examples of simulated home range areas (km^2^) of 0.64 (b-c), 2.54 (d), 10.18 (e) and 91.61 (f). Home range centres (red dots), home range areas (red circles).

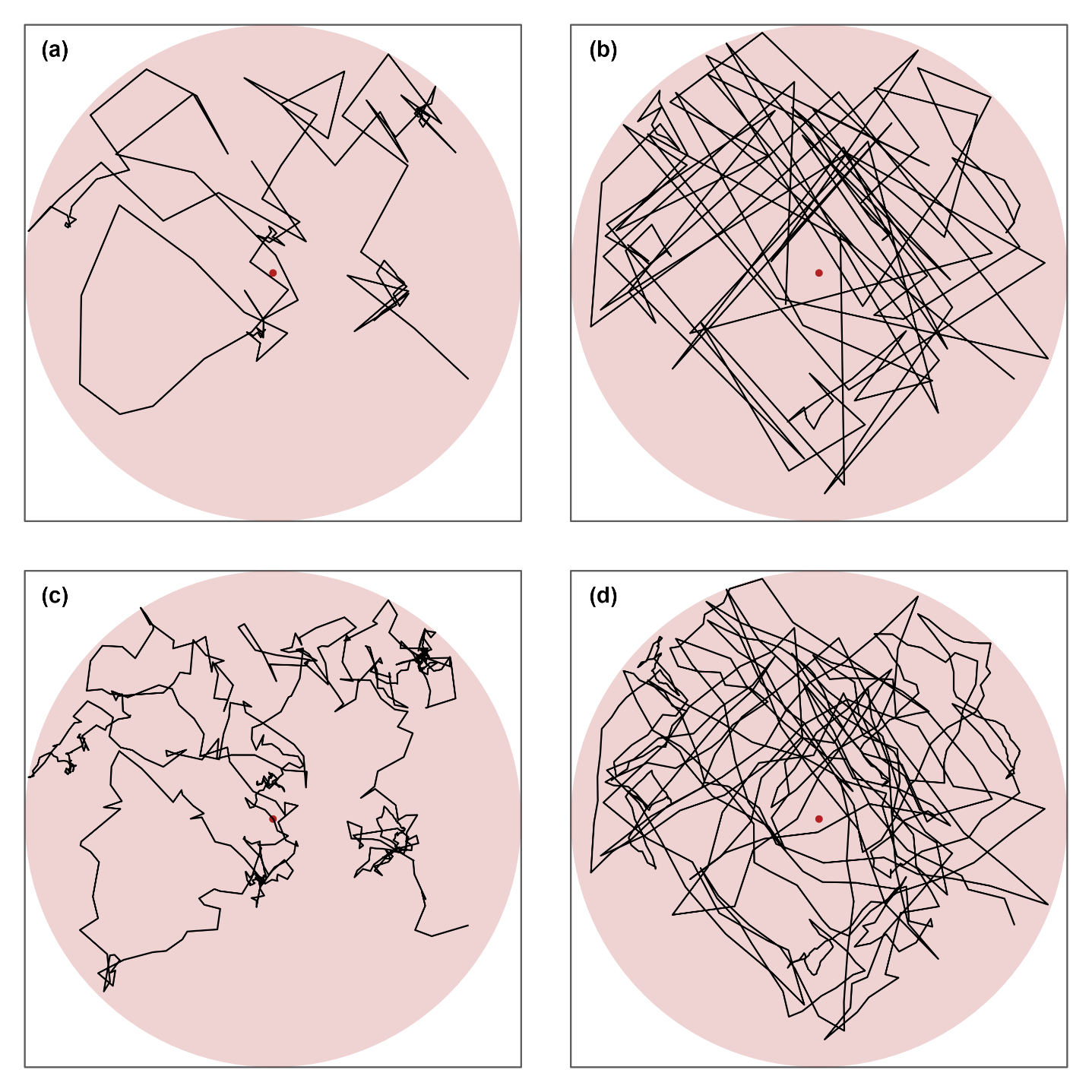

**Figure A.2:** An example of two simulated movement trajectories (only five days are displayed for visual clarity) within a given home range area, before (a-b) and after (c-d) the addition of Brownian motion. (a) One displacement per hour, slow movements; (b) one displacement per hour, fast movements; (c) six displacements per hour (every 10 min.), slow movements; and (d) six displacements per hour (every 10 min.), fast movements. Home range centers (red dots), home range areas (red circles) and movement trajectories (full and dotted black lines).

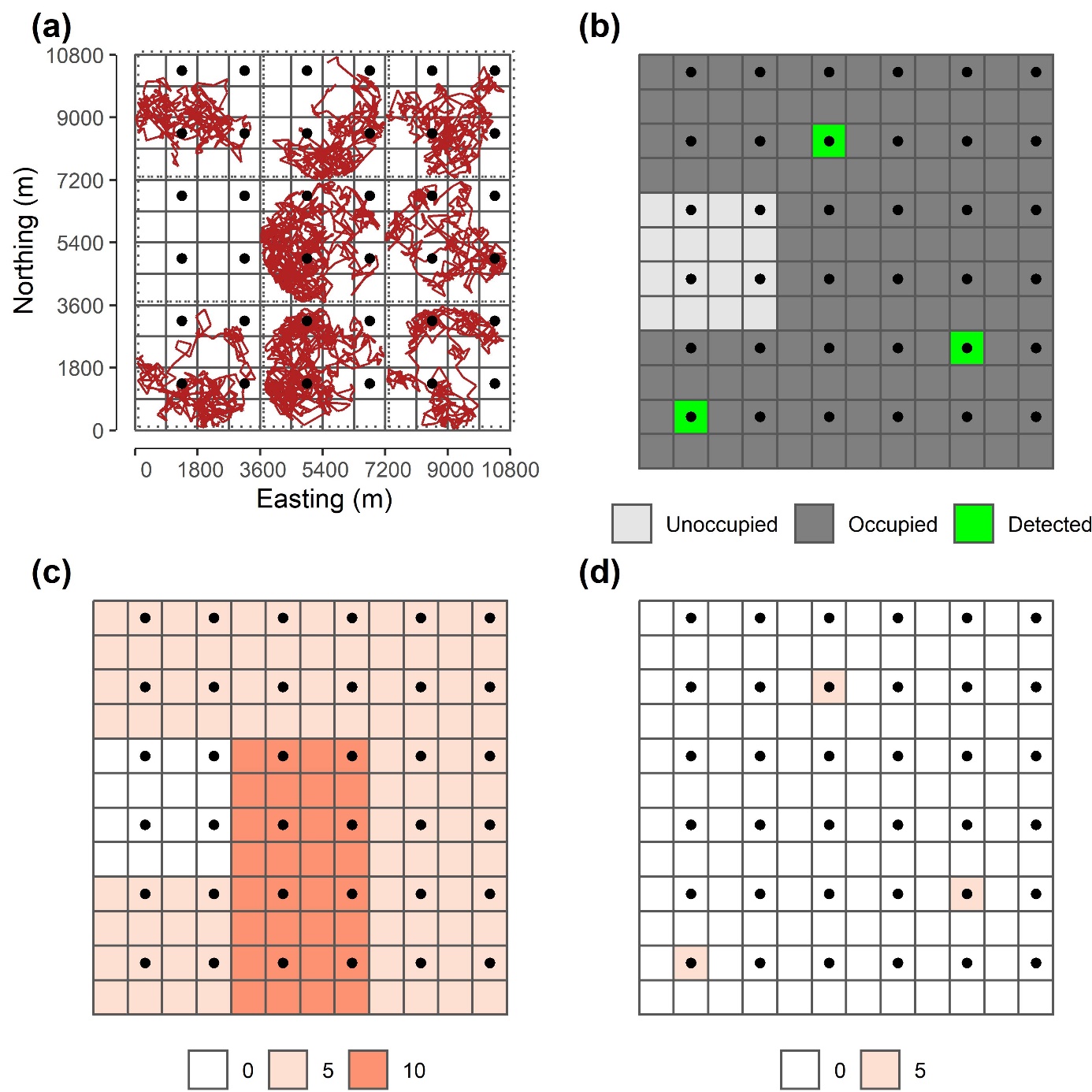

**Figure A.3:** A summary of the data generating mechanism. (a) Movement trajectories (red) within the state space. (b) The asymptotic occupancy status of the sampling grid cells: unoccupied (light grey), occupied (grey) or occupied with a detection (green). (c) The asymptotic abundance per sampling grid cells. (d) The number of unmarked counts per sampling grid cell. Bounding boxes for group-specific home ranges are represented by dotted lines, the sampling grid cells by solid lines and camera deployments by black dots.
