## Supplementary material for "Simulation-based assessment of the performance of hierarchical abundance estimators for camera trap surveys": Appendix B_Stanmodels.docx

Appendix B. Parametrization, prior specification, model fitting, goodness-of-fit and MCMC convergence of hierarchical abundance models fit in Stan

For all the hierarchical abundance models (HMs) considered in our simulation study, we choose vaguely informative priors for detection parameters $\theta_{det}$ and site-occupancy (in zero-inflated Poisson mixtures) $\psi$, i.e., $Uniform\left( 0, 1 \right)$, and a weakly informative $Cauchy\left( 0, 10 \right)$ prior for the site-abundance $\lambda$. The Stan codes for each HM fitted in our study can be found on LINK. These models were fitted using four parallel MCMC chains, with 2000 iterations, which included 1000 iterations that were discarded as burn-in iterations. In total our simulation study consists of 900 different scenarios, for which we fit 40 models each (*i.e.,* one for each simulation replicate). To make it computationally efficient to fit all these models, we use the high-performance computing (HPC) clusters provided by the VSC (Flemish Supercomputer Center). We run our models in parallel using 8 threads, with 36 nodes per thread. Moreover, we break up the likelihood statement in partial sums and split the calculations over 8 threads per chain using Stan’s *reduce_sum* built-in function. To evaluate the predictive performance and the goodness-of-fit of competing HMs, we calculate respectively the Leave-one-out expected log predictive density (LOO ELPD; **Figure B.1**; **Table B.1**), and Bayesian *P*-values (**Figure B.3**; **Table B.1**) in the generated quantities block of Stan, as explained in section 2 of this appendix. Finally, we assess the model convergence by the $\hat{R}$-statistic and the sampling efficiency in the bulk and tail of the posterior distribution by the effective sample size (ESS; **Figure B.5**-**8**).

### 1 Goodness-of-fit and predictive performance

To assess goodness-of-fit for competing HMs in our study, we calculate Bayesian *P*-values.

For each posterior sample, $s=1, 2,..., 4000$, we generated data $y_{ij}^{(s)}$ and retrieve expected values $E_{ij}^{(s)}$, then calculated

$$T^{(s)}=\sum_{i} \sum_{j} \frac{{(y_{ij}^{(s)}-E_{ij}^{(s)})}^{2}}{E_{ij}^{(s)}}$$

The proportion of samples for which $T^{\left( s \right)}$ exceeds the value of the same statistic, computed with actual data $y_{ij}$ rather than $y_{ij}^{\left( s \right)}$, is the Bayesian *P*-value $P_{B}$ (Hjort et al., 2006; Link et al., 2018). Note that expected values $E_{ij}^{\left( s \right)}$, needed to calculate $T^{(s)}$, depend on the HM,

$$E_{ij}^{\left( s \right)}=\left\{ \begin{aligned} {1-\left( 1-p^{\left( s \right)} \right)}^{N_{i}^{\left( s \right)}} if Bernoulli-Poisson mixture \\ N_{i}^{\left( s \right)}\cdot p^{\left( s \right)} if Binomial-Poisson mixture \\ \left( N_{i}^{\left( s \right)} \right)^{2}\cdot p^{\left( s \right)} if Poisson-Poisson mixture \end{aligned} \right.$$

In addition to $P_{B}$’s, we also calculated leave-one-out (LOO) expected log predictive densities (Vehtari et al., 2017) to assess the predictive performance of competing HMs. The LOO, $\chi^{2}$-squared discrepancy between replicated data $y_{ij}^{(s)}$ and expected values $E_{ij}^{(s)}$, $T^{(s)}$, the $P_{B}$’s and the proportions of simulations with $P_{B}\leq0.05$ are visualized in **Figures B.1**-**B.4**. For a tabular representation, consult **Table B.1**.

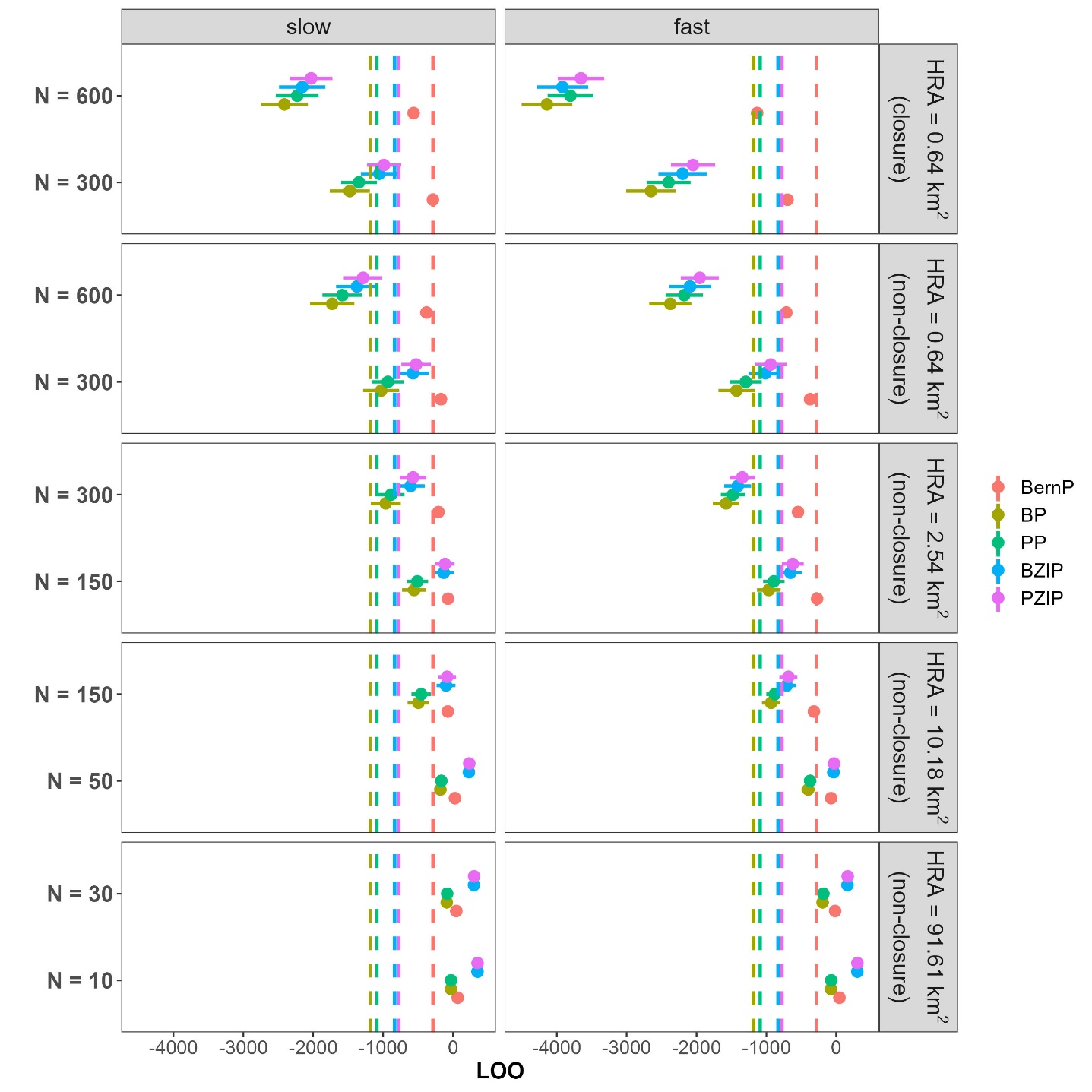

**Figure B.1.** LOO for all combinations of model used (BernP, BP, PP, BZIP, PZIP), movement speed (slow, fast), population size N (10, 30, 50, 150, 300, 600) and home range area in km^2^ (0.64, 2.54, 10.18, 91.61). Dots and horizontal lines represent means and standard errors for LOO in each scenario, while the vertical dashed line indicates the average LOO overall.

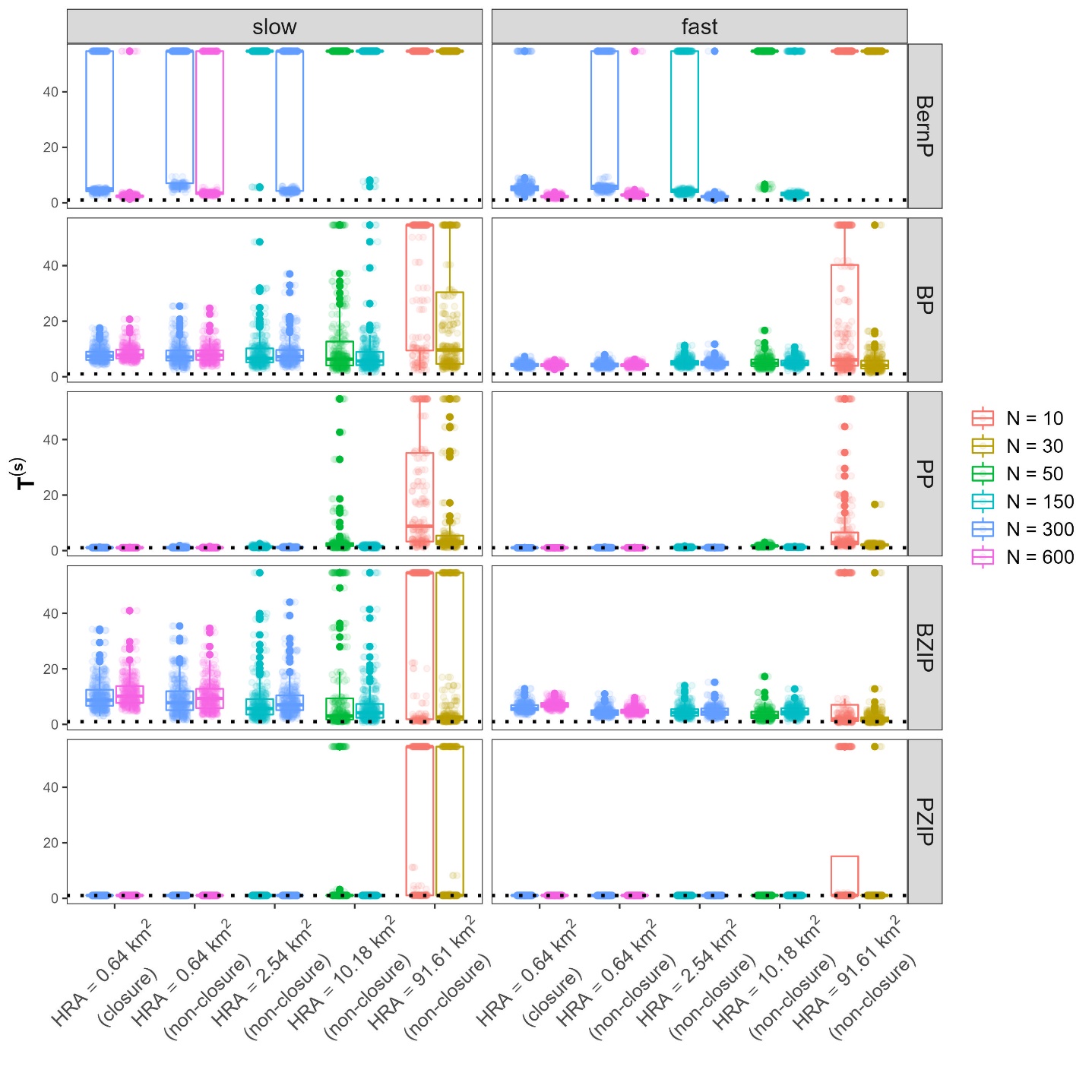

**Figure B.2.** $\chi^{2}$-discrepancy between replicated data and expected values, for all combinations of model used (BernP, BP, PP, BZIP, PZIP), movement speed (slow, fast), population size N (10, 30, 50, 150, 300, 600) and home range area in km^2^ (0.64, 2.54, 10.18, 91.61). For graphical clarity, values exceeding the 85^th^ percentile were truncated.

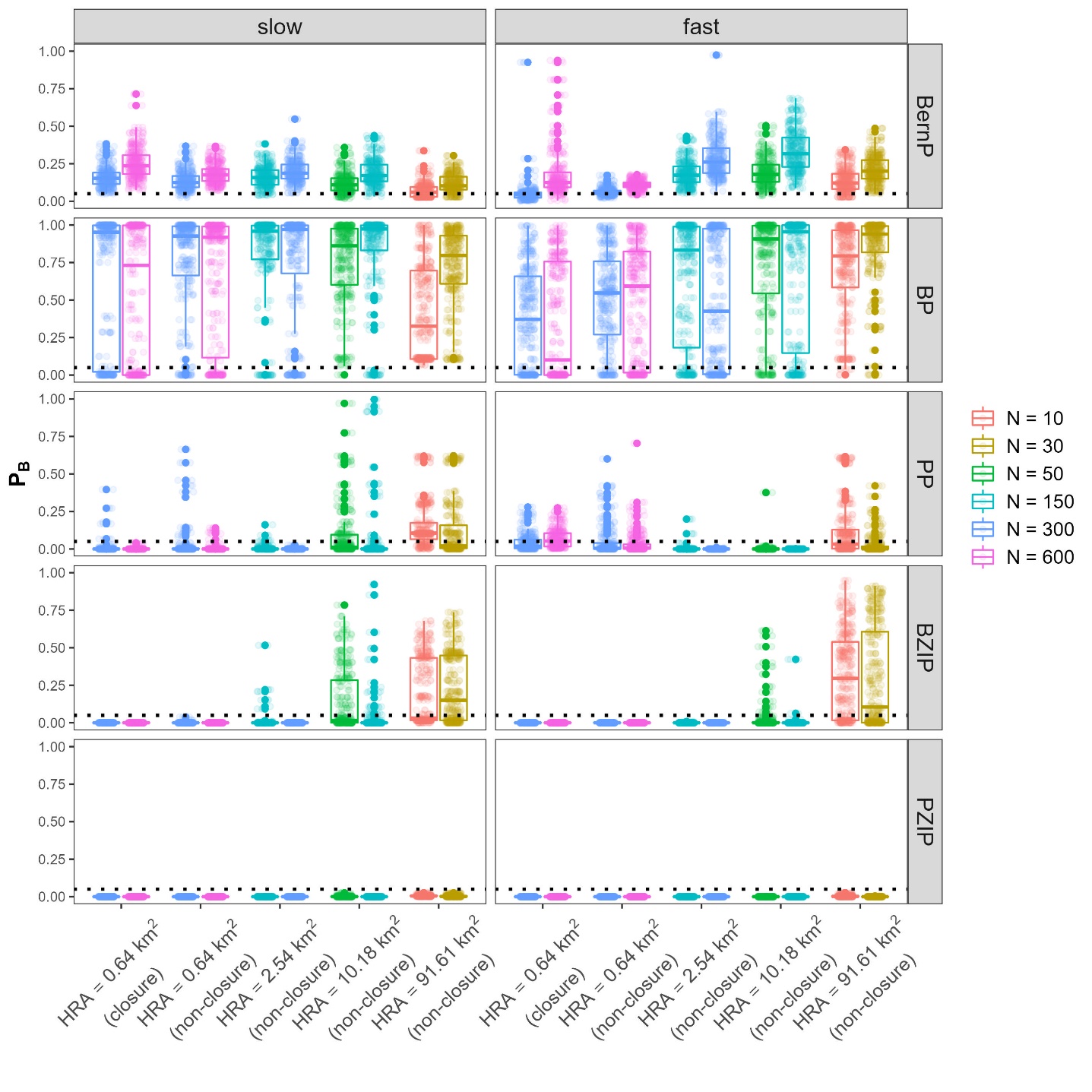

**Figure B.3.** Bayesian *P*-values $P_{B}$ for all combinations of model used (BernP, BP, PP, BZIP, PZIP), movement speed (slow, fast), population size N (10, 30, 50, 150, 300, 600) and home range area in km^2^ (0.64, 2.54, 10.18, 91.61).

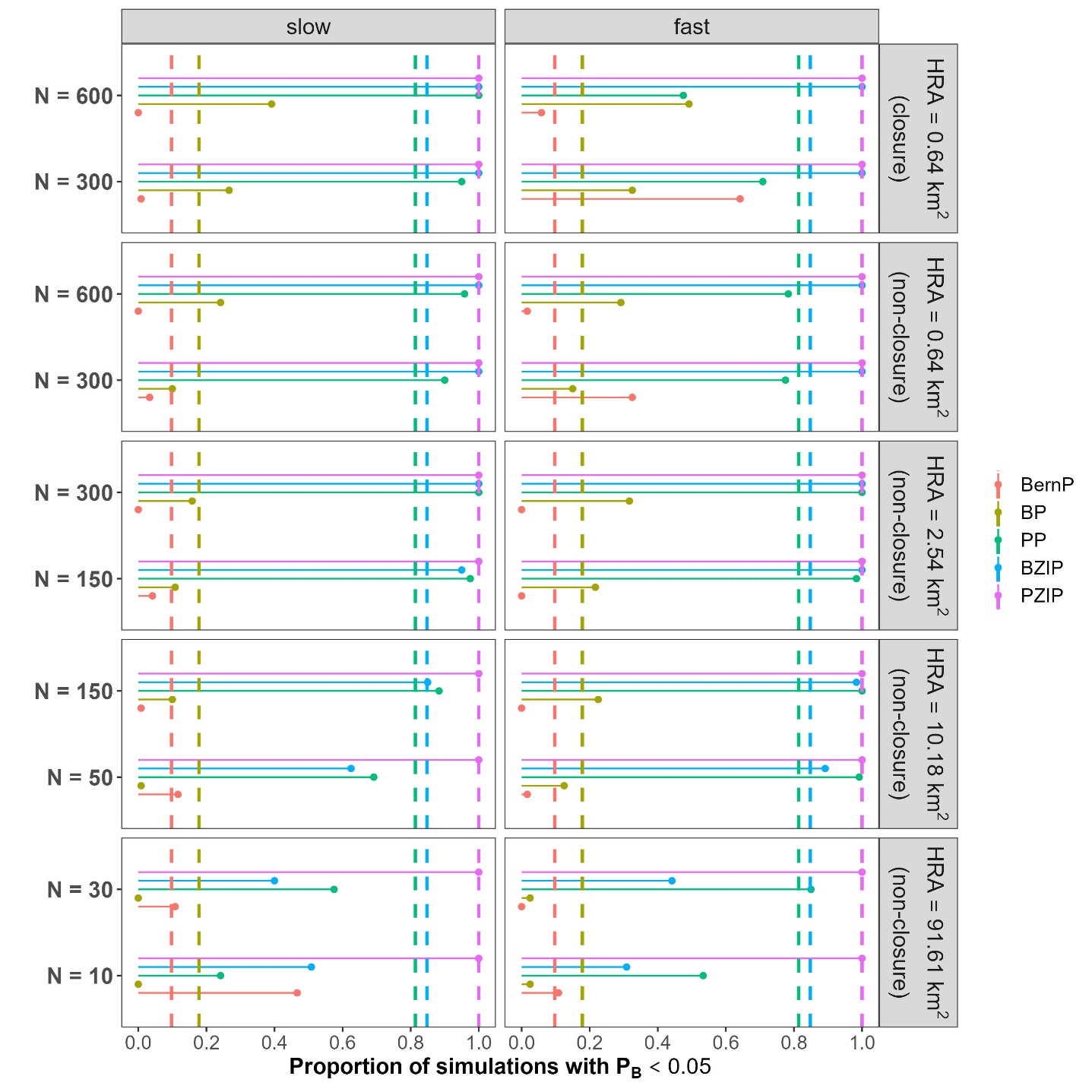

**Figure B.4.** Proportion of simulations with Bayesian *P*-values $P_{B}$ < 0.05 for all combinations of model used (BernP, BP, PP, BZIP, PZIP), movement speed (slow, fast), population size N (10, 30, 50, 150, 300, 600) and home range area in km^2^ (0.64, 2.54, 10.18, 91.61). The dots represent proportions in each scenario, while the vertical dashed line indicates the average proportion of $P_{B}$ < 0.05 overall.

|  |  |  | **Proportion of simulations with** $\boldsymbol{P}_{\boldsymbol{b}}\boldsymbol{\leq0.05}$ | | | | | **LOO** | | | | |
| --- | --- | --- | --- | --- | --- | --- | --- | --- | --- | --- | --- | --- |
| **HRA (km^2^)** | **N** | **Speed** | **BernP** | **BP** | **PP** | **BZIP** | **PZIP** | **BernP** | **BP** | **PP** | **BZIP** | **PZIP** |
| 0.64  (closure) | 600 | slow | **0.00** | 0.39 | 1.00 | 1.00 | 1.00 | **-562.94** | -2412.06 | -2227.95 | -2157.01 | -2029.02 |
|  |  | fast | **0.06** | 0.49 | 0.48 | 1.00 | 1.00 | **-1132.02** | -4140.43 | -3805.21 | -3918.00 | -3653.89 |
|  | 300 | slow | **0.01** | 0.27 | 0.95 | 1.00 | 1.00 | **-286.52** | -1476.55 | -1344.88 | -1052.93 | -986.69 |
|  |  | fast | **0.64** | 0.33 | 0.71 | 1.00 | 1.00 | **-698.73** | -2651.39 | -2400.12 | -2199.71 | -2049.84 |
| 0.64 | 300 | slow | **0.00** | 0.24 | 0.96 | 1.00 | 1.00 | **-378.85** | -1727.99 | -1582.76 | -1372.73 | -1286.46 |
|  |  | fast | **0.02** | 0.29 | 0.78 | 1.00 | 1.00 | **-714.87** | -2375.19 | -2176.03 | -2093.71 | -1953.08 |
|  | 150 | slow | **0.03** | 0.10 | 0.90 | 1.00 | 1.00 | **-171.18** | -1026.69 | -931.21 | -571.81 | -527.65 |
|  |  | fast | **0.33** | 0.15 | 0.78 | 1.00 | 1.00 | **-375.74** | -1427.39 | -1295.21 | -1012.70 | -938.29 |
| 2.54 | 150 | slow | **0.00** | 0.16 | 1.00 | 1.00 | 1.00 | **-206.96** | -961.56 | -887.49 | -605.78 | -570.76 |
|  |  | fast | **0.00** | 0.32 | 1.00 | 1.00 | 1.00 | **-547.58** | -1576.55 | -1479.00 | -1410.31 | -1345.44 |
|  | 50 | slow | **0.04** | 0.11 | 0.98 | 0.95 | 1.00 | **-70.75** | -556.51 | -509.29 | -131.13 | -113.91 |
|  |  | fast | **0.00** | 0.22 | 0.98 | 1.00 | 1.00 | **-274.87** | -965.91 | -893.37 | -656.80 | -622.63 |
| 10.18 | 150 | slow | **0.01** | 0.10 | 0.88 | 0.85 | 1.00 | **-73.61** | -494.52 | -455.34 | -98.92 | -83.48 |
|  |  | fast | **0.00** | 0.23 | 1.00 | 0.98 | 1.00 | **-320.01** | -932.09 | -880.45 | -709.54 | -685.71 |
|  | 50 | slow | 0.12 | **0.01** | 0.69 | 0.63 | 1.00 | 28.10 | -180.62 | -166.43 | 228.01 | **232.80** |
|  |  | fast | **0.02** | 0.13 | 0.99 | 0.89 | 1.00 | -74.61 | -401.80 | -374.86 | -37.97 | **-31.10** |
| 91.61 | 30 | slow | 0.11 | **0.00** | 0.58 | 0.40 | 1.00 | 48.29 | -88.66 | -82.63 | 300.76 | **302.75** |
|  |  | fast | **0.00** | 0.03 | 0.85 | 0.44 | 1.00 | -17.07 | -195.91 | -184.93 | 160.23 | **163.02** |
|  | 10 | slow | 0.47 | **0.00** | 0.24 | 0.51 | 1.00 | 68.54 | -28.90 | -27.23 | 351.36 | **351.95** |
|  |  | fast | 0.11 | **0.03** | 0.53 | 0.31 | 1.00 | 45.44 | -77.19 | -72.82 | 301.42 | **302.42** |
| **Average** | | | **0.10** | 0.18 | 0.81 | 0.85 | 1.00 | **-285.80** | -1184.89 | -1088.86 | -834.36 | -776.25 |

**Table B.1:** Summary table for the goodness-of-fit and predictive performance of BernP, BP, PP, BZIP and PZIP. LOO, leave-one-out expected log predictive density; $P_{b}$, Bayesian *P*-value. Note that the highest proportion of simulations with $P_{b}\leq0.05$, and the highest LOOs are indicated in bold.

### 2 MCMC convergence and sampling efficiency Figure B.5. Proportion of MCMC chains that ended with a divergence for all combinations of model used (BernP, BP, PP, BZIP, PZIP) movement speed (slow, fast), population size N (10, 30, 50, 150, 300, 600) and home range area in km^2^ (0.64, 2.54, 10.18, 91.61).

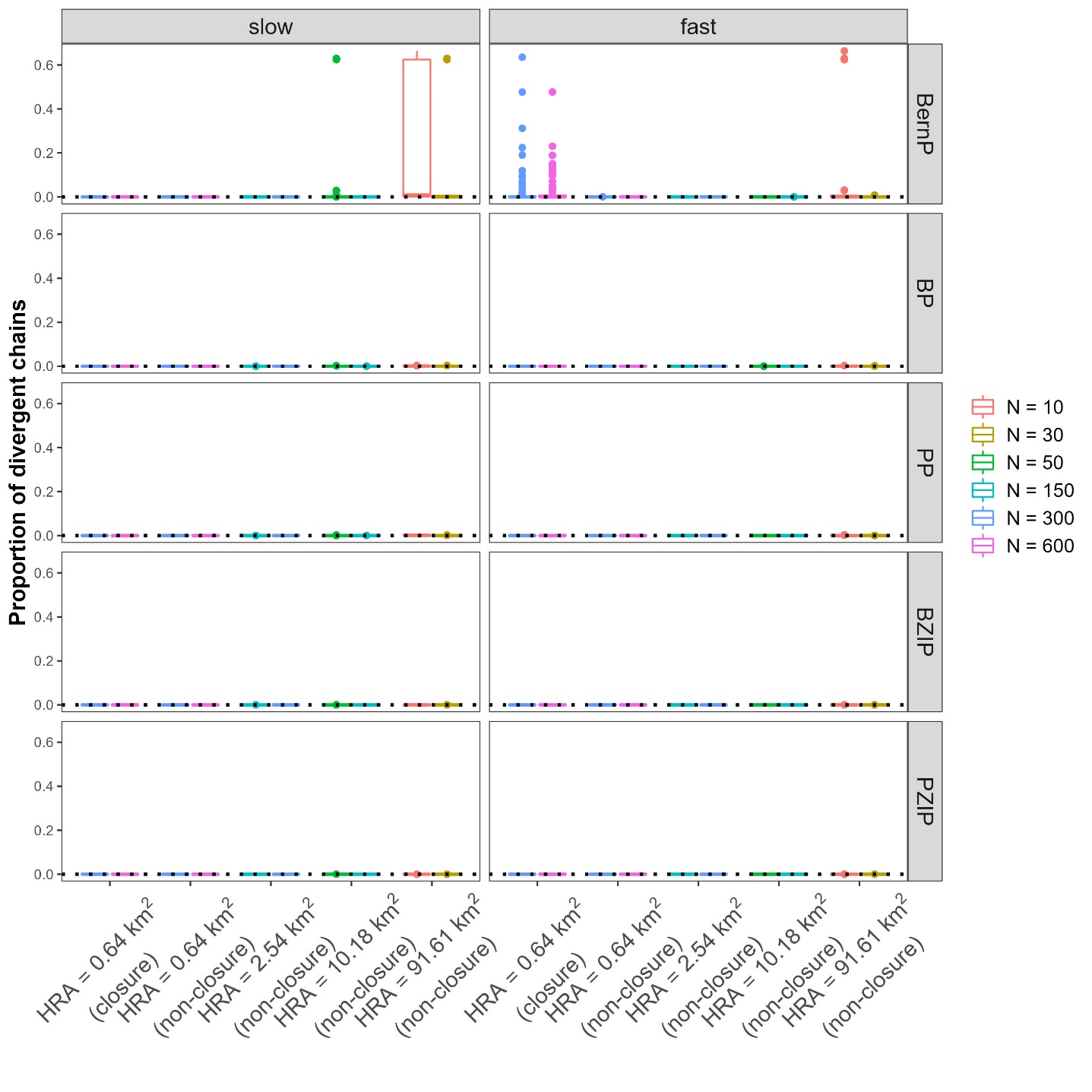

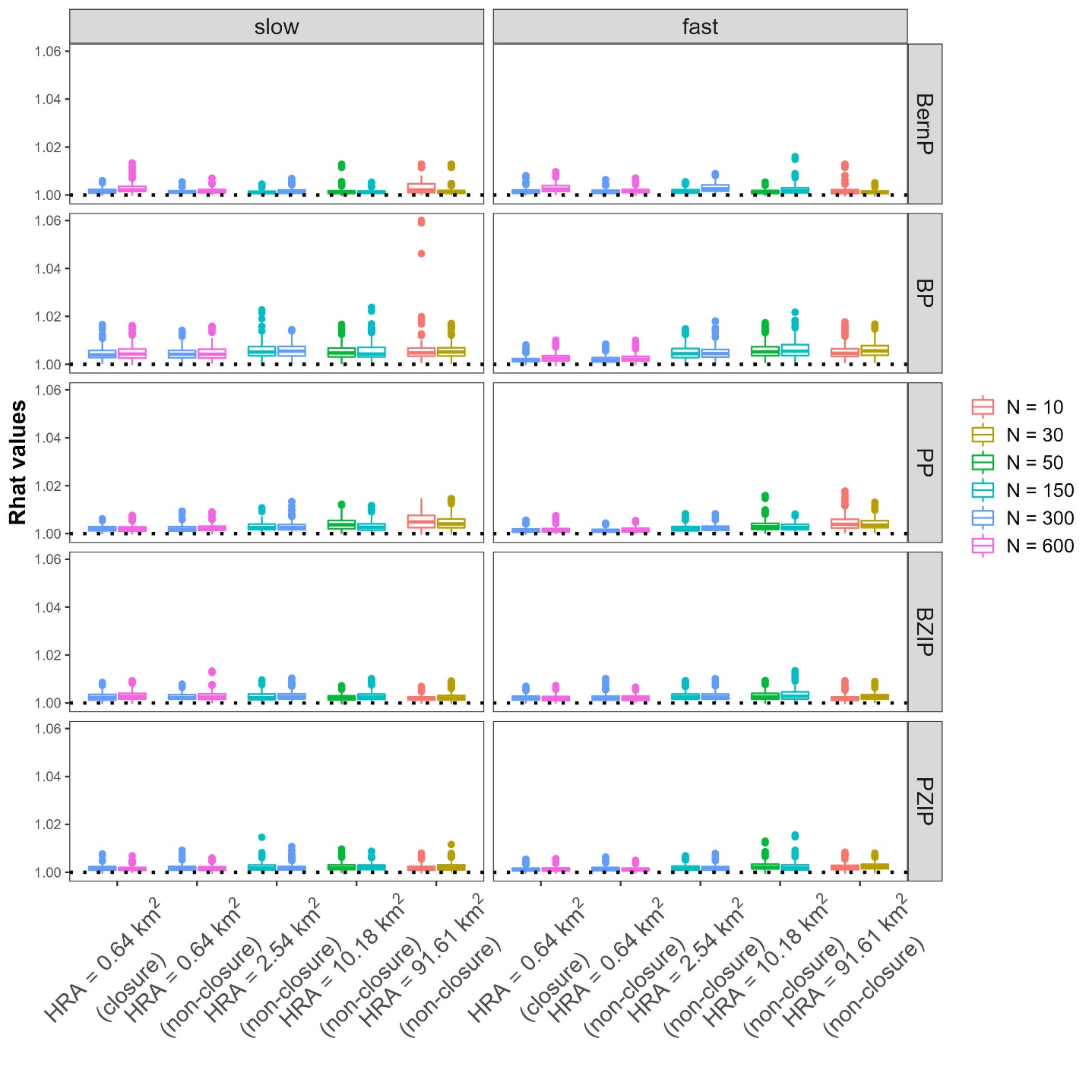

**Figure B.6.** $\hat{R}$-statistic for all combinations of model used (BernP, BP, PP, BZIP, PZIP), movement speed (slow, fast), population size N (10, 30, 50, 150, 300, 600) and home range area in km^2^ (0.64, 2.54, 10.18, 91.61).

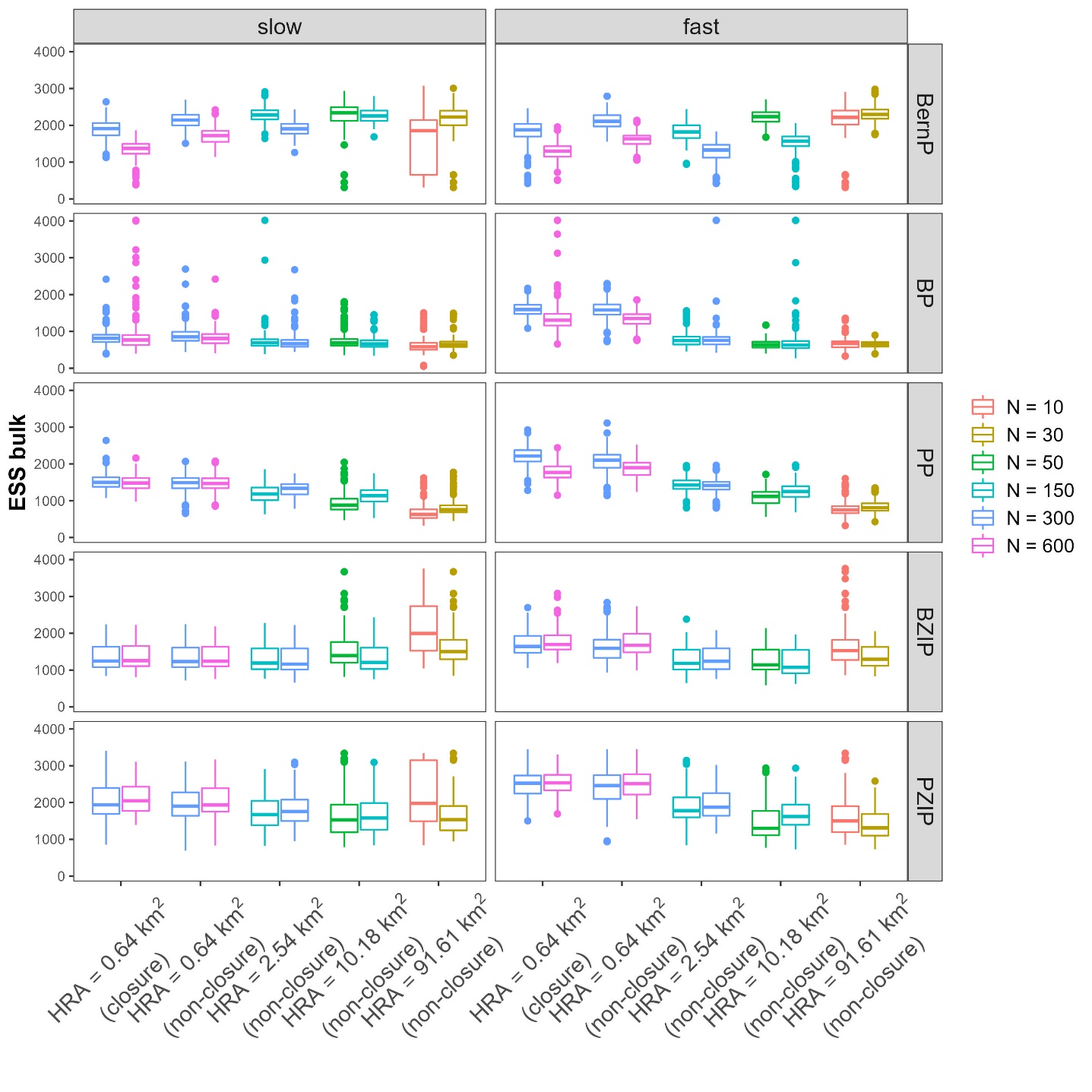

**Figure B.7.** ESS bulk for all combinations of model used (BernP, BP, PP, BZIP, PZIP), movement speed (slow, fast), population size N (10, 30, 50, 150, 300, 600) and home range area in km^2^ (0.64, 2.54, 10.18, 91.61).

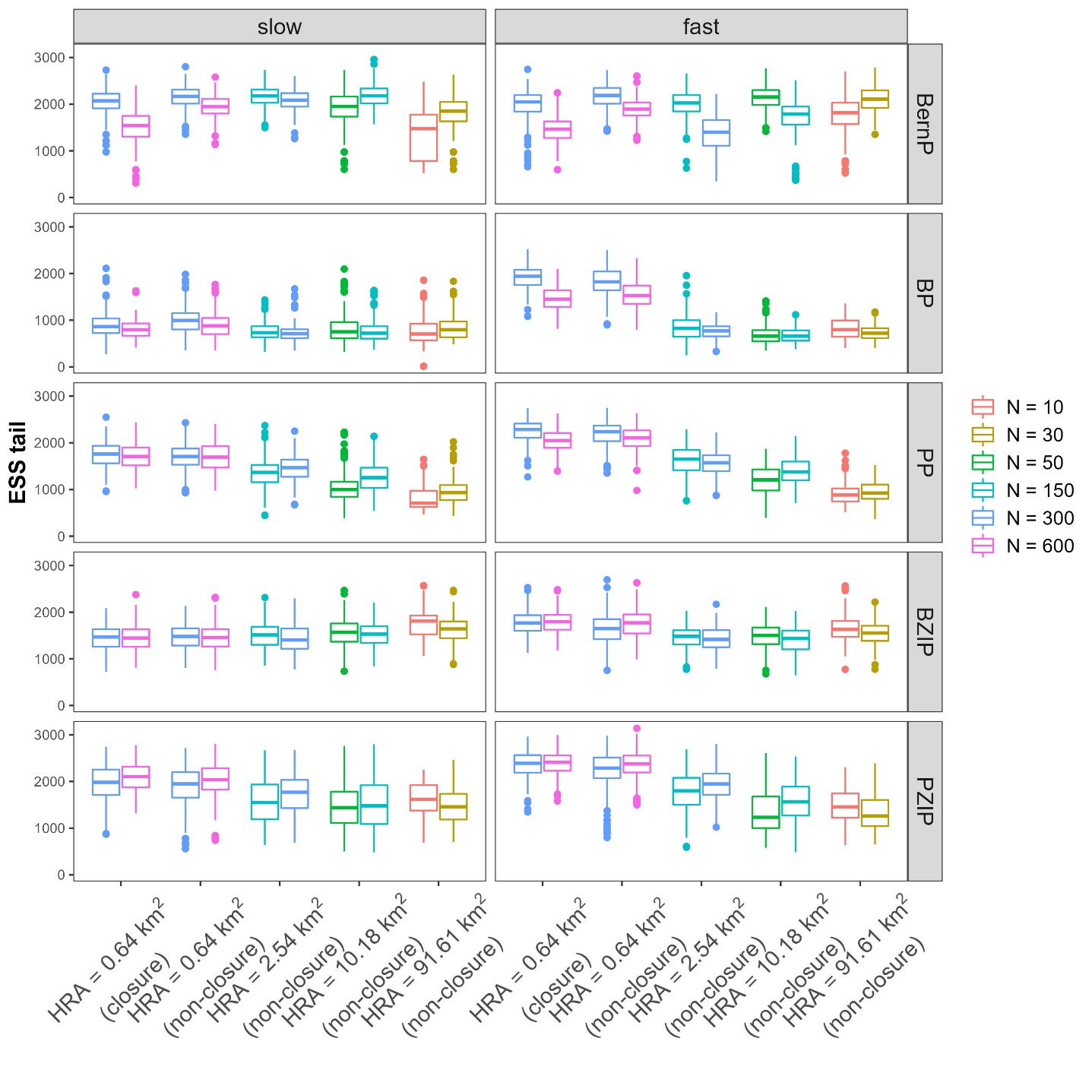

**Figure B.8.** ESS tail for all combinations of model used (BernP, BP, PP, BZIP, PZIP), movement speed (slow, fast), population size N (10, 30, 50, 150, 300, 600) and home range area in km^2^ (0.64, 2.54, 10.18, 91.61).
