## Supplementary material for "Simulation-based assessment of the performance of hierarchical abundance estimators for camera trap surveys": Appendix C_vis_all_scenarios.docx

Appendix C. Visualisations and summary tables for all simulation scenarios

In the main paper, the direction of bias in estimated detection parameters $\theta_{det}$, abundances $\lambda$ and site use frequencies $\lambda_{use}$ are shown for all of the models (**Figure 3**). However, the scenarios with home range areas of 91.61 km^2^ were omitted form this figure. Here, we display similar figures, including the scenarios omitted in the main paper (**Figures C.1-4**). Moreover, this appendix contains summary tables that display the 95% credible interval (CI) coverages and root mean square errors for BernP, BP and PP models (**Tables C.1-C.3**). Finally, it also contains summary tables that display the proportion of simulations where |Relative Bias| ≤ 0.5, the 95% credible interval (CI) coverages and root mean square errors for BZIP and PZIP models (**Tables C.4-C.6**).

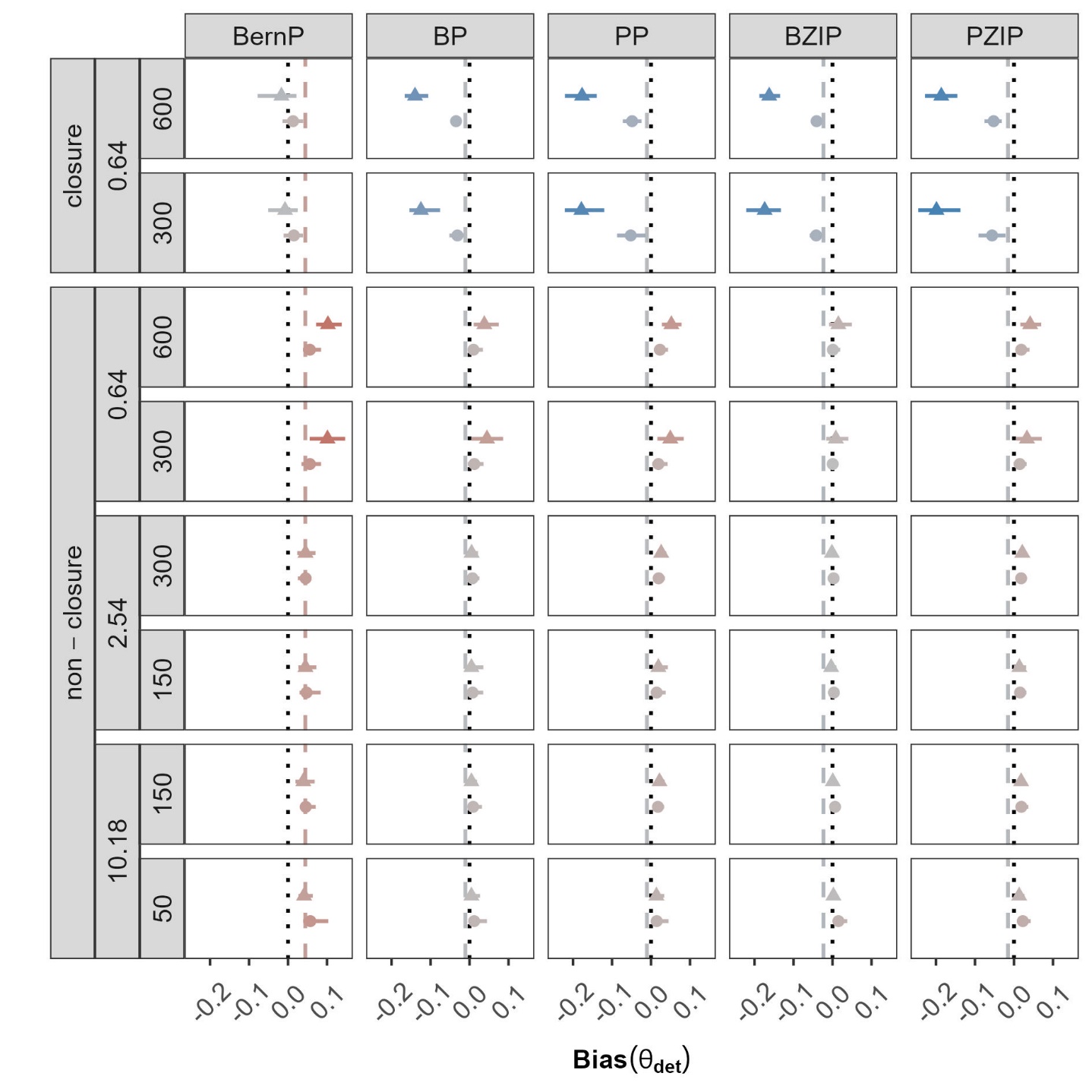

**Figure C.1:** Mean bias (dots/ triangles), together with 2.5% and 97.5% quantiles (solid lines) in the estimated detection parameters $\theta_{det}$. Results are displayed for all combinations of population size (N = 10, 30, 50, 150, 300, 600), speed of movement (slow: triangles, fast: dots), home range area (HRA) in km^2^ (0.64, 2.54, 10.18 and 91.81), geographical closure (closure, non-closure) and hierarchical model fitted (BernP, BP, PP, BZIP and PZIP). Line of equality (dotted line). Average bias for each HM (dashed line).

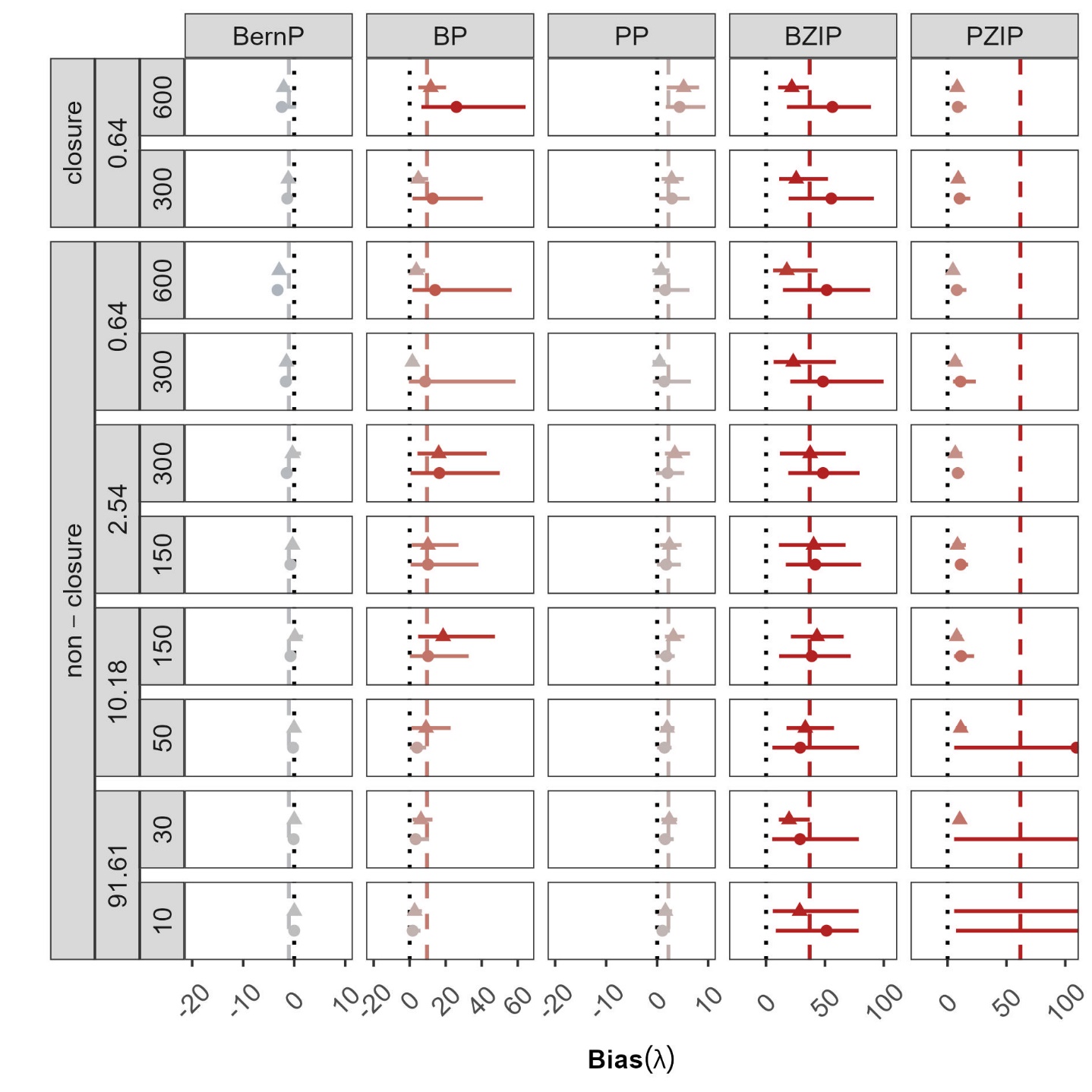

**Figure C.2:** Mean bias (dots/ triangles), together with 2.5% and 97.5% quantiles (solid lines) in the estimated abundances $\lambda$. Results are displayed for all combinations of population size (N = 10, 30, 50, 150, 300, 600), speed of movement (slow: triangles, fast: dots), home range area (HRA) in km^2^ (0.64, 2.54, 10.18 and 91.81), geographical closure (closure, non-closure) and hierarchical model fitted (BernP, BP, PP, BZIP and PZIP). Line of equality (dotted line). Average bias for each HM (dashed line). For visual clarity, x-scales are different for the subpanels.

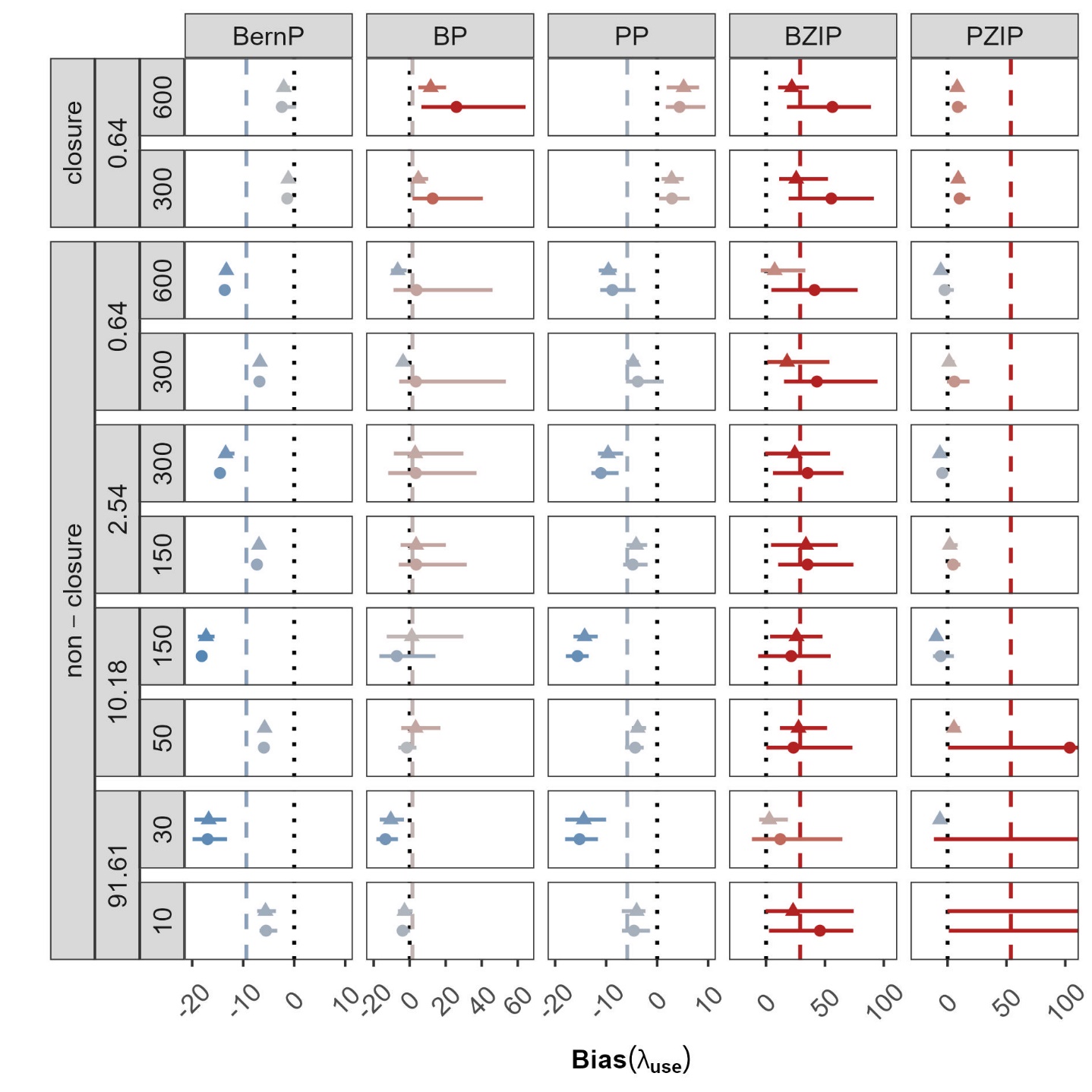

**Figure C.3:** Mean bias (dots/ triangles), together with 2.5% and 97.5% quantiles (solid lines) in the estimated site use frequencies $\lambda_{use}$. Results are displayed for all combinations of population size (N = 10, 30, 50, 150, 300, 600), speed of movement (slow: triangles, fast: dots), home range area (HRA) in km^2^ (0.64, 2.54, 10.18 and 91.81), geographical closure (closure, non-closure) and hierarchical model fitted (BernP, BP, PP, BZIP and PZIP). Line of equality (dotted line). Average bias for each HM (dashed line). For visual clarity, x-scales are different for the subpanels

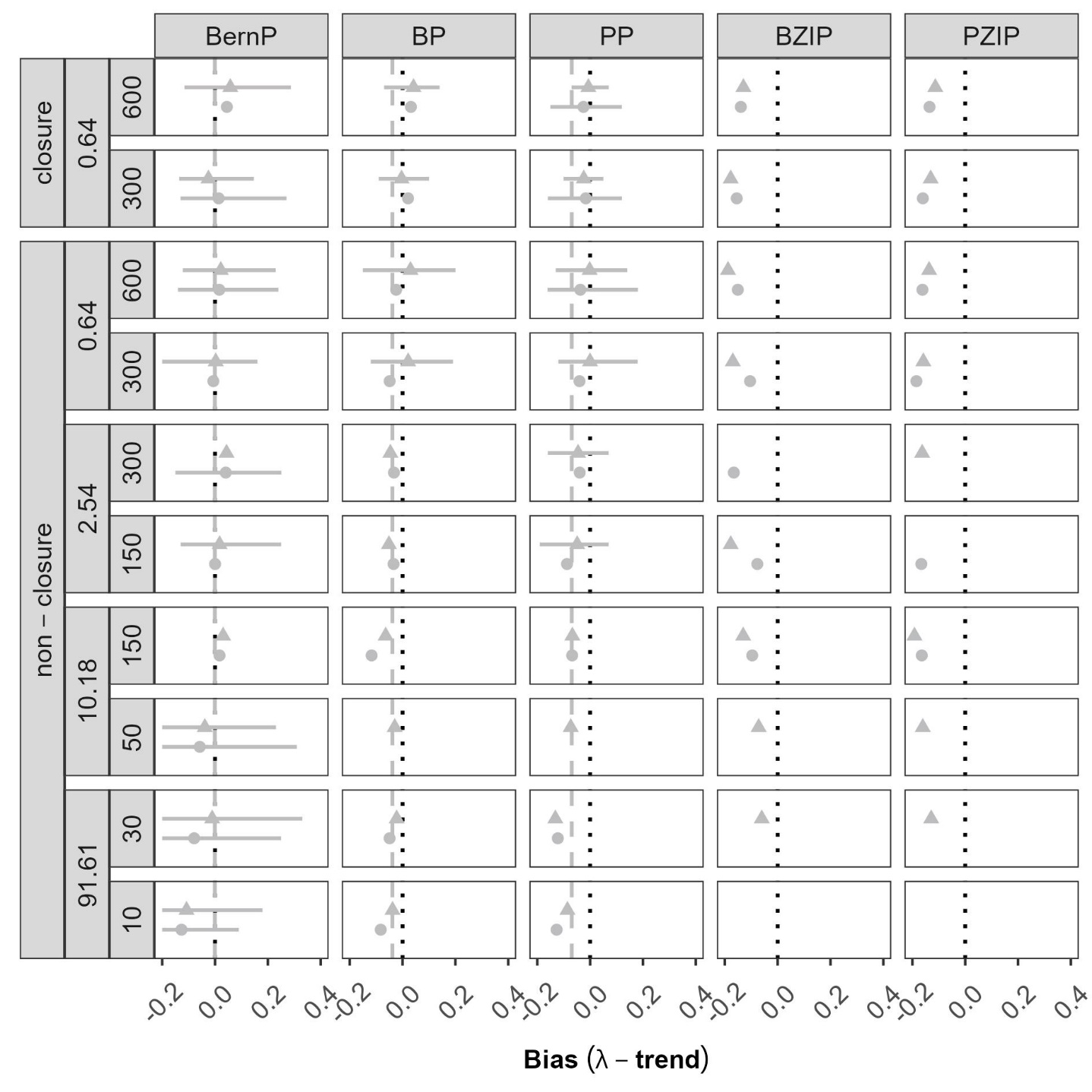

**Figure C.4:** Mean bias (dots/ triangles), together with 2.5% and 97.5% quantiles (solid lines) in the estimated trend bias in abundances $\lambda$. Results are displayed for all combinations of population size (N = 10, 30, 50, 150, 300, 600), speed of movement (slow: triangles, fast: dots), home range area (HRA) in km^2^ (0.64, 2.54, 10.18 and 91.81), geographical closure (closure, non-closure) and hierarchical model fitted (BernP, BP, PP, BZIP and PZIP). Line of equality (dotted line). Average bias for each HM (dashed line).

**Table C.1:** Summary table for estimator quality of detection parameters $\theta_{det}$ obtained from three Bayesian hierarchical models (BernP, BP, PP). Cells display the 95% CI coverage and the root mean square error based on forty simulation replicates (note that the highest CI coverages and lowest root mean square errors in each scenario are indicated in bold).

|  |  |  | **95% CI coverage** | | | **Root mean square error** | | |
| --- | --- | --- | --- | --- | --- | --- | --- | --- |
| **HRA (km^2^)** | **N** | **Speed** | **BernP** | **BP** | **PP** | **BernP** | **BP** | **PP** |
| 0.64  (closure) | 600 | slow | **75.00** | 2.50 | 0.00 | **0.02** | 0.04 | 0.05 |
|  |  | fast | **85.00** | 0.00 | 0.00 | **0.03** | 0.14 | 0.18 |
|  | 300 | slow | **70.00** | 2.50 | 0.00 | **0.02** | 0.03 | 0.06 |
|  |  | fast | **92.50** | 0.00 | 0.00 | **0.03** | 0.13 | 0.18 |
| 0.64 | 600 | slow | 0.00 | **40.00** | 0.00 | 0.06 | **0.01** | 0.03 |
|  |  | fast | 0.00 | **2.50** | 0.00 | 0.10 | **0.04** | 0.05 |
|  | 300 | slow | 0.00 | **37.50** | 12.50 | 0.06 | **0.02** | **0.02** |
|  |  | fast | 0.00 | **5.00** | 0.00 | 0.10 | **0.05** | **0.05** |
| 2.54 | 300 | slow | 0.00 | **52.50** | 0.00 | 0.05 | **0.01** | 0.02 |
|  |  | fast | 2.50 | **52.50** | 0.00 | 0.05 | **0.01** | 0.03 |
|  | 150 | slow | 0.00 | **67.50** | 27.50 | 0.05 | **0.01** | 0.02 |
|  |  | fast | 0.00 | **67.50** | 5.00 | 0.05 | **0.01** | 0.02 |
| 10.18 | 150 | slow | 0.00 | **42.50** | 2.50 | 0.05 | **0.01** | 0.02 |
|  |  | fast | 5.00 | **67.50** | 0.00 | 0.04 | **0.01** | 0.02 |
|  | 50 | slow | 0.00 | **65.00** | 32.50 | 0.07 | 0.13 | 0.13 |
|  |  | fast | 0.00 | **80.00** | 22.50 | **0.04** | **0.01** | 0.02 |
| 91.61 | 30 | slow | 0.00 | **12.50** | 2.50 | **0.08** | 0.17 | 0.17 |
|  |  | fast | 0.00 | **40.00** | 7.50 | 0.04 | **0.01** | **0.01** |
|  | 10 | slow | 0.00 | **22.50** | 20.00 | **0.12** | 0.30 | 0.30 |
|  |  | fast | 0.00 | **52.50** | 45.00 | **0.09** | 0.19 | 0.19 |
| **Median** | | | 0.00 | **40.00** | 1.25 | 0.05 | **0.03** | 0.04 |

**Table C.2:** Summary table for estimator quality of abundances $\lambda$ obtained from three Bayesian hierarchical models (BernP, BP, PP). Cells display the 95% CI coverage and the root mean square error based on forty simulation replicates (note that the highest CI coverages and lowest root mean square errors in each scenario are indicated in bold).

|  |  |  | **95% CI coverage** | | | **Root mean square error** | | |
| --- | --- | --- | --- | --- | --- | --- | --- | --- |
| **HRA (km^2^)** | **N** | **Speed** | **BernP** | **BP** | **PP** | **BernP** | **BP** | **PP** |
| 0.64  (closure) | 600 | slow | **5.00** | 0.00 | 0.00 | **2.62** | 31.32 | 4.93 |
|  |  | fast | **7.50** | 0.00 | 0.00 | **2.20** | 12.46 | 5.48 |
|  | 300 | slow | **5.00** | 0.00 | **5.00** | **1.38** | 18.52 | 3.40 |
|  |  | fast | **2.50** | **2.50** | **2.50** | **1.23** | 5.27 | 3.10 |
| 0.64 | 600 | slow | 0.00 | 0.00 | **50.00** | 3.28 | 20.63 | **2.64** |
|  |  | fast | 0.00 | 12.50 | **37.50** | 2.95 | 4.40 | **1.27** |
|  | 300 | slow | 0.00 | 12.50 | **40.00** | **1.65** | 15.53 | 2.28 |
|  |  | fast | 0.00 | 32.50 | **52.50** | 1.52 | 2.13 | **0.86** |
| 2.54 | 300 | slow | 0.00 | 5.00 | **30.00** | **1.48** | 22.22 | 2.61 |
|  |  | fast | **60.00** | 0.00 | 0.00 | **0.80** | 19.32 | 3.73 |
|  | 150 | slow | 0.00 | 7.50 | **25.00** | **0.75** | 14.78 | 2.24 |
|  |  | fast | **40.00** | 0.00 | 2.50 | **0.37** | 12.19 | 2.67 |
| 10.18 | 150 | slow | 0.00 | 15.00 | **27.50** | **0.73** | 13.88 | 2.53 |
|  |  | fast | **80.00** | 0.00 | 0.00 | **0.76** | 22.36 | 3.39 |
|  | 50 | slow | 5.00 | 42.50 | **52.50** | **0.24** | 5.54 | 1.79 |
|  |  | fast | **92.50** | 2.50 | 2.50 | **0.10** | 10.83 | 2.07 |
| 91.61 | 30 | slow | 27.50 | 57.50 | **60.00** | **0.13** | 4.39 | 1.81 |
|  |  | fast | **95.00** | 2.50 | 2.50 | **0.08** | 7.02 | 2.57 |
|  | 10 | slow | **100.00** | 67.50 | 72.50 | **0.03** | 2.53 | 1.31 |
|  |  | fast | **97.50** | 40.00 | 40.00 | **0.04** | 3.56 | 1.86 |
| **Median** | | | 5.00 | 3.75 | **26.25** | **0.78** | 12.33 | 2.55 |

**Table C.3:** Summary table for estimator quality of site use frequencies $\lambda_{use}$ obtained from three Bayesian hierarchical models (BernP, BP, PP). Cells display the 95% CI coverage and the root mean square error based on forty simulation replicates (note that the highest CI coverages and lowest root mean square errors in each scenario are indicated in bold).

|  |  |  | **95% CI coverage** | | | **Root mean square error** | | |
| --- | --- | --- | --- | --- | --- | --- | --- | --- |
| **HRA (km^2^)** | **N** | **Speed** | **BernP** | **BP** | **PP** | **BernP** | **BP** | **PP** |
| 0.64  (closure) | 600 | slow | **5.00** | 0.00 | 0.00 | **2.62** | 31.32 | 4.93 |
|  |  | fast | **7.50** | 0.00 | 0.00 | **2.20** | 12.46 | 5.48 |
|  | 300 | slow | **5.00** | 0.00 | **5.00** | **1.38** | 18.52 | 3.40 |
|  |  | fast | **2.50** | **2.50** | **2.50** | **1.23** | 5.27 | 3.10 |
| 0.64 | 600 | slow | 0.00 | **55.00** | 2.50 | 13.63 | 15.45 | **9.05** |
|  |  | fast | 0.00 | **10.00** | 0.00 | 13.31 | 7.11 | **9.61** |
|  | 300 | slow | 0.00 | **52.50** | 10.00 | 6.81 | 13.36 | **4.24** |
|  |  | fast | 0.00 | **7.50** | 0.00 | 6.71 | 3.99 | **4.76** |
| 2.54 | 300 | slow | 0.00 | **60.00** | 0.00 | 14.56 | 15.20 | **11.16** |
|  |  | fast | 0.00 | **65.00** | 0.00 | 13.46 | 11.06 | **9.74** |
|  | 150 | slow | 0.00 | **72.50** | 12.50 | 7.31 | 11.10 | **4.99** |
|  |  | fast | 0.00 | **60.00** | 2.50 | 6.91 | 7.64 | **4.26** |
| 10.18 | 150 | slow | 0.00 | **55.00** | 0.00 | 18.15 | **11.86** | 15.76 |
|  |  | fast | 0.00 | **70.00** | 0.00 | 17.30 | **12.58** | 14.30 |
|  | 50 | slow | 0.00 | **67.50** | 30.00 | 6.01 | **4.23** | 4.50 |
|  |  | fast | 0.00 | **80.00** | 10.00 | 5.82 | 6.81 | **3.92** |
| 91.61 | 30 | slow | 0.00 | **25.00** | 0.00 | 16.99 | **13.93** | 15.34 |
|  |  | fast | 0.00 | **65.00** | 2.50 | 16.85 | **11.04** | 14.53 |
|  | 10 | slow | 0.00 | **35.00** | 27.50 | 5.67 | **4.47** | 4.74 |
|  |  | fast | 0.00 | **65.00** | 55.00 | 5.71 | **3.72** | 4.22 |
| **Median** | | | 0.00 | **55.00** | 2.50 | 6.86 | 11.08 | **4.96** |

|  |  |  | **\|Relative Bias\| ≤ 0.5** | | **95% CI coverage** | | **Root mean square error** | |
| --- | --- | --- | --- | --- | --- | --- | --- | --- |
| **HRA (km^2^)** | **N** | **Speed** | **BZIP** | **PZIP** | **BZIP** | **PZIP** | **BZIP** | **PZIP** |
| 0.64  (closure) | 600 | slow | 0.03 | **0.25** | 0.00 | 0.00 | **0.04** | 0.05 |
|  |  | fast | 0.00 | 0.00 | 0.00 | 0.00 | **0.16** | 0.19 |
|  | 300 | slow | 0.03 | **0.13** | 0.00 | 0.00 | **0.04** | 0.06 |
|  |  | fast | 0.00 | 0.00 | 0.00 | 0.00 | **0.18** | 0.20 |
| 0.64 | 600 | slow | **0.85** | 0.13 | **72.50** | 7.50 | **0.01** | 0.02 |
|  |  | fast | **0.38** | 0.00 | **30.00** | 0.00 | **0.02** | 0.04 |
|  | 300 | slow | **0.78** | 0.25 | **75.00** | 30.00 | **0.00** | 0.02 |
|  |  | fast | **0.53** | 0.20 | **52.50** | 15.00 | **0.02** | 0.04 |
| 2.54 | 300 | slow | **0.45** | 0.00 | **82.50** | 0.00 | **0.01** | 0.02 |
|  |  | fast | **0.68** | 0.00 | **62.50** | 0.00 | **0.01** | 0.02 |
|  | 150 | slow | **0.35** | 0.10 | **77.50** | 22.50 | **0.01** | 0.02 |
|  |  | fast | **0.65** | 0.25 | **60.00** | 37.50 | **0.01** | 0.02 |
| 10.18 | 150 | slow | 0.00 | 0.00 | **10.00** | 0.00 | **0.01** | 0.02 |
|  |  | fast | **0.78** | 0.00 | **90.00** | 2.50 | **0.00** | 0.02 |
|  | 50 | slow | 0.00 | 0.00 | **7.50** | **7.50** | **0.16** | **0.16** |
|  |  | fast | **0.58** | 0.03 | **90.00** | 40.00 | **0.00** | 0.01 |
| 91.61 | 30 | slow | 0.00 | 0.00 | 0.00 | 0.00 | **0.21** | 0.21 |
|  |  | fast | 0.00 | 0.00 | 0.00 | 0.00 | **0.01** | 0.02 |
|  | 10 | slow | 0.00 | 0.00 | 0.00 | 0.00 | 0.39 | **0.38** |
|  |  | fast | 0.00 | 0.00 | 0.00 | 0.00 | **0.24** | **0.24** |
| **Median** | | | **0.19** | 0.00 | **20.00** | 0.00 | **0.01** | 0.03 |

**Table C.4:** Summary table for estimator quality of detection parameters $\theta_{det}$ obtained from two Bayesian hierarchical models (BZIP, PZIP). Cells display the proportion of simulation replicates that satisfy |Relative Bias| ≤ 0.5, the 95% CI coverage and the root mean square error based on forty simulation replicates (note that the highest proportions, CI coverages and lowest root mean square errors in each scenario are indicated in bold).

|  |  |  | **\|Relative Bias\| ≤ 0.5** | | **95% CI coverage** | | **Root mean square error** | |
| --- | --- | --- | --- | --- | --- | --- | --- | --- |
| **HRA (km^2^)** | **N** | **Speed** | **BZIP** | **PZIP** | **BZIP** | **PZIP** | **BZIP** | **PZIP** |
| 0.64  (closure) | 600 | slow | 0.00 | 0.00 | 0.00 | 0.00 | 59.60 | **9.19** |
|  |  | fast | 0.00 | 0.00 | 0.00 | 0.00 | 22.83 | **8.44** |
|  | 300 | slow | 0.00 | 0.00 | 0.00 | 0.00 | 58.52 | **10.95** |
|  |  | fast | 0.00 | 0.00 | 0.00 | 0.00 | 28.16 | **9.44** |
| 0.64 | 600 | slow | 0.00 | 0.00 | 0.00 | 0.00 | 55.31 | **8.77** |
|  |  | fast | 0.00 | **0.10** | 0.00 | 0.00 | 21.32 | **4.86** |
|  | 300 | slow | 0.00 | 0.00 | 0.00 | 0.00 | 52.66 | **11.96** |
|  |  | fast | 0.00 | 0.00 | 0.00 | 0.00 | 27.76 | **6.86** |
| 2.54 | 300 | slow | 0.00 | 0.00 | 0.00 | 0.00 | 52.02 | **9.22** |
|  |  | fast | 0.00 | 0.00 | 0.00 | 0.00 | 40.88 | **6.88** |
|  | 150 | slow | 0.00 | 0.00 | 0.00 | 0.00 | 45.50 | **11.73** |
|  |  | fast | 0.00 | 0.00 | 0.00 | 0.00 | 42.89 | **8.90** |
| 10.18 | 150 | slow | 0.00 | 0.00 | 0.00 | 0.00 | 43.39 | **12.69** |
|  |  | fast | 0.00 | 0.00 | 0.00 | 0.00 | 44.86 | **8.00** |
|  | 50 | slow | 0.00 | 0.00 | 0.00 | 0.00 | **38.15** | 313.23 |
|  |  | fast | 0.00 | 0.00 | 0.00 | 0.00 | 35.31 | **11.47** |
| 91.61 | 30 | slow | 0.00 | 0.00 | 0.00 | 0.00 | **39.90** | 414.26 |
|  |  | fast | 0.00 | 0.00 | 0.00 | 0.00 | **21.29** | **10.42** |
|  | 10 | slow | 0.00 | 0.00 | 0.00 | 0.00 | 60.76 | 750.76 |
|  |  | fast | 0.00 | 0.00 | 0.00 | 0.00 | **39.85** | 469.69 |
| **Median** | | | 0.00 | 0.00 | 0.00 | 0.00 | 41.89 | **9.93** |

**Table C.5:** Summary table for estimator quality of abundances $\lambda$ obtained from two Bayesian hierarchical models (BZIP, PZIP). Cells display the proportion of simulation replicates that satisfy |Relative Bias| ≤ 0.5, the 95% CI coverage and the root mean square error based on forty simulation replicates (note that the highest proportions, CI coverages and lowest root mean square errors in each scenario are indicated in bold).

|  |  |  | **\|Relative Bias\| ≤ 0.5** | | **95% CI coverage** | | **Root mean square error** | |
| --- | --- | --- | --- | --- | --- | --- | --- | --- |
| **HRA (km^2^)** | **N** | **Speed** | **BZIP** | **PZIP** | **BZIP** | **PZIP** | **BZIP** | **PZIP** |
| 0.64  (closure) | 600 | slow | 0.00 | 0.00 | 0.00 | 0.00 | 59.60 | **9.19** |
|  |  | fast | 0.00 | 0.00 | 0.00 | 0.00 | 22.83 | **8.44** |
|  | 300 | slow | 0.00 | 0.00 | 0.00 | 0.00 | 58.52 | **10.95** |
|  |  | fast | 0.00 | 0.00 | 0.00 | 0.00 | 28.16 | **9.44** |
| 0.64 | 600 | slow | 0.05 | **0.95** | 2.50 | **35.00** | 45.79 | **4.58** |
|  |  | fast | 0.65 | **0.83** | **37.50** | 2.50 | 14.01 | **6.06** |
|  | 300 | slow | 0.00 | **0.40** | 2.50 | **50.00** | 47.97 | **7.52** |
|  |  | fast | 0.13 | **0.83** | 12.50 | **57.50** | 23.61 | **2.59** |
| 2.54 | 300 | slow | 0.08 | **0.80** | 25.00 | **60.00** | 40.09 | **5.51** |
|  |  | fast | 0.18 | **0.70** | **27.50** | 12.50 | 29.36 | **6.88** |
|  | 150 | slow | 0.00 | **0.43** | 10.00 | **72.50** | 39.52 | **5.79** |
|  |  | fast | 0.03 | **0.83** | 5.00 | **85.00** | 36.76 | **3.33** |
| 10.18 | 150 | slow | 0.25 | **0.80** | 62.50 | **80.00** | 28.97 | **8.12** |
|  |  | fast | 0.08 | **0.43** | **42.50** | 17.50 | 28.46 | **9.78** |
|  | 50 | slow | 0.10 | **0.28** | 57.50 | **90.00** | **33.95** | 311.16 |
|  |  | fast | 0.03 | **0.15** | 12.50 | **80.00** | 29.91 | **5.97** |
| 91.61 | 30 | slow | 0.40 | **0.43** | 92.50 | **95.00** | **30.28** | 407.35 |
|  |  | fast | 0.78 | **0.95** | **97.50** | **97.50** | 8.57 | **6.73** |
|  | 10 | slow | 0.10 | **0.20** | 90.00 | **97.50** | **56.09** | 746.48 |
|  |  | fast | 0.18 | **0.28** | 80.00 | **100.00** | **36.03** | 466.98 |
| **Median** | | | 0.08 | **0.43** | 18.75 | **58.75** | 32.11 | **7.82** |

**Table C.6:** Summary table for estimator quality of site use frequencies $\lambda_{use}$ obtained from two Bayesian hierarchical models (BZIP, PZIP). Cells display the proportion of simulation replicates that satisfy |Relative Bias| ≤ 0.5, the 95% CI coverage and the root mean square error based on forty simulation replicates (note that the highest proportions, CI coverages and lowest root mean square errors in each scenario are indicated in bold).
